# MHC II-expressing bone marrow megakaryocytes are noncanonical antigen presenting cells and activate CD4^+^ T cells ex vivo

**DOI:** 10.1101/2025.11.21.689743

**Authors:** Virginia Camacho, Karen G. Wang, Pavel Hanč, Estelle Carminita, Isabelle C. Becker, Dong H. Lee, Mahmoud A. Bassal, Marcelo Falchetti, Jaxaira Maggi, Maria Barrachina, Ulrich von Andrian, Deepa Gautam, Chen Weng, Vijay G. Sankaran, Montserrat Carrascal, Joseph E. Italiano, Kellie R. Machlus

## Abstract

While professional antigen-presenting cells drive adaptive immunity, atypical cell types can fulfill this role in the bone marrow. Megakaryocytes (MKs) are canonically recognized for platelet production, but recent studies indicate functional heterogeneity and immune potential. We found that ∼20% of bone marrow MKs express Major Histocompatibility Complex (MHC) II and co-stimulatory receptors CD80, CD86, CD40, and CD83. These MKs process and present antigen to activate T cells ex vivo in an MHC II-dependent manner. MK/T cell interactions induced TGF-β1 secretion and promoted induced Treg differentiation. Prior stimulation of MKs with LPS or Poly I:C was associated with modest Th1-associated CD4+ T cell responses, including IFN-γ and TNF-α production, without robust Th17 differentiation. Immunopeptidomics of the murine MK MHC II receptor confirmed occupancy by exogenous peptides, suggesting in vivo functionality. Using a murine model with MK-targeted deletion of MHC II *(Pf4-MHC*Δ/Δ), we observed altered TLR signaling and reduced bone marrow TGF-β1. Together, these findings identify MHC II+ MKs as noncanonical antigen-presenting cells with the potential to modulate CD4 T cell responses as part of the immune regulation of the bone marrow niche.

## INTRODUCTION

Megakaryocytes (**MKs**) are large (50–100 µm) cells derived from hematopoietic stem cells (**HSCs**) in the bone marrow (1). MKs are canonically recognized for the ability to generate platelets, circulating cells essential for blood clotting and wound healing. However, recent single cell analyses of human and murine bone marrow MKs have revealed transcriptional heterogeneity within this population and demonstrate that MKs have spatially and functionally specialized roles within this niche (2–4). As such, the range of functions attributed to MKs has significantly expanded, evolving from cells that make platelets to cells that play roles in HSC support, bone marrow niche regulation, and the instruction of adaptive immunity. Sequencing studies have classified a subset of MKs as ‘immune MKs’ based on the enrichment of genes associated with antigen presentation and toll-like receptor (**TLR**) signaling (2, 4). These and other studies suggest that MKs may communicate with T cells through direct contact involving major histocompatibility complex (**MHC**) class I, II, and co-stimulatory molecules, supporting the notion that a subset of MKs can shape adaptive immune responses within the bone marrow. Nonetheless, there has been insufficient evidence confirming a functional antigen presentation capacity of MKs in the bone marrow and lack of comprehensive characterization of these cells or their significance.

The presentation of peptides by antigen presenting cells (**APCs**) is a key driver in the initiation and regulation of adaptive immunity. MHC class I is found on all nucleated cells and presents antigenic peptides to CD8+ T cells. Conversely, constitutive MHC class II expression is restricted to ‘professional APCs’. While dendritic cells are regarded as the typical professional APC of the immune system, dendritic cells are not solely responsible for priming CD4+ T cell responses. Atypical cell types can execute this role in tissues such as the bone marrow with a unique immunological profile (5–7). In the bone marrow, HSCs can process and present antigen via MHC II (8). This safeguards HSCs from exhaustion and prevents malignant transformation. Furthermore, in HSCs, MHC II expression increases with age, and these MHC II+ aged HSCs are found more proximal to regulatory T cells (**Tregs**). Specifically, the interaction between MHC II+ HSCs and Tregs helps aged HSCs maintain long-term survival by creating an environment that limits the effects of inflammation and immune attack (9). This expands on other studies demonstrating that bone marrow Tregs directly and indirectly support the maintenance and function of HSCs, as well as the stromal microenvironment via IL-10 (9, 10). Together, these studies support the notion of CD4+ T cell crosstalk with hematopoietic cells in the bone marrow to regulate hematopoiesis.

Notably, MKs also have an established role in regulating hematopoiesis. The spatial positioning of MKs next to HSCs not only modifies HSC activity but also influences lineage output; HSCs adjacent to MKs exhibit a greater platelet- and myeloid-biased differentiation profile compared to HSCs in other marrow regions, such as the arteriolar niche. Furthermore, MK depletion can reprogram these myeloid-biased HSCs toward a more lineage-balanced state (11). This is partially executed via cytokine release; MKs produce TGF-β1 and PF4, which help maintain HSC quiescence in vivo (11). MKs also support HSC niche remodeling, including osteoblast expansion, following radiation injury and promotion of HSC proliferation under stress conditions by producing FGF1 and IGF-1 (12, 13). Thus, MKs are established regulators of the bone marrow microenvironment. However, it is unknown if hematopoietic regulation is driven by specific subsets of bone marrow MKs or connected to the immune functions of MKs.

In this manuscript, we characterize MHC II-expressing MKs in the bone marrow. We confirmed the presence of MHC II on a subset of bone marrow MKs in both mice and humans. Notably, MHC II was absent in platelets, even when platelets were activated, suggesting that while MKs express this molecule, they do not readily transfer it to platelets. We observed that stimulation of MKs with TLR agonists such as poly I:C and LPS enhanced the expression of MHC II and co-stimulatory molecules, indicating functional TLR responses. Bone marrow MKs harbored the ability to activate CD4+ T cells in an MHC II-dependent manner ex vivo and induce Treg differentiation. Further, transcriptomic profiling of MHC II+ MKs revealed upregulated interferon responses and enhanced phagocytic capacity. Finally, MK-specific deletion of MHC II *(Pf4-MHC*^Δ/Δ^ mice) altered MK responsiveness to TLR agonists in vivo, reduced bone marrow TGF-β1 production, and altered stem and progenitor cell responses to TLR stimulation in the bone marrow. Overall, this work highlights a noncanonical role for MHC II+ MKs as immunologic regulators in the bone marrow.

## RESULTS

### MHC II is expressed throughout the megakaryocyte lineage

To characterize MHC II expression in the myelo-erythroid progenitor populations in the bone marrow, we utilized a murine MHC II–EGFP knock-in model, in which green fluorescent protein (**EGFP**) is fused to the MHC II beta chain and expression is driven by endogenous transcriptional activity (14, 15). This model provided a tool for identifying MHC II expressing cells while limiting potential artifacts associated with antibody-based labeling approaches. Using MHC II–EGFP knock-in mice, we compared MHC II across various stem and progenitor populations in the bone marrow. These data revealed that MHC II expression was prevalent in the MK lineage; 17.7% ± 5.07 (mean ± SD) of mature MKs (Lineage-CD41+ CD42d+) expressed MHC II (**Fig 1A-B**). Further, approximately 39% (± 15.74) of MkPs (CD150+ CD41+ Lineage-Sca-1-c-kit+) expressed MHC II (**Fig 1A-B** and **S1A-B**) and MK-biased HSCs were also enriched in MHC II expression (57.4 ± 21.6, **Fig 1A, C**). MK-erythroid progenitors expressed low amounts of MHC II, with expression decreasing along the erythroid lineage and reappearing in granulocyte-monocyte progenitors (**Fig 1C, S1A**). MHC II expression was also visualized using high-dimensional flow cytometry, displaying its co-expression with CD41+ MKs and MK progenitor cells (**Fig 1D**). Given the fragility and large size of native MKs, we sought to further substantiate MHC II expression on MKs by visualization *in situ*. Imaging of femoral cryosections confirmed endogenous MHC II expression in a subset of MKs through EGFP and CD41+ co-expression (**Fig 1E**). These findings were corroborated with live cell imaging of native, bone marrow MKs where cell surface MHC II was observed in a subpopulation of large, polyploid MKs (**Fig 1F**). Notably, we also observed expression of MHC II in splenic MKs, with an average expression level of 33.37 ± 4.88 (**Fig S1B**), as previously reported (16, 17). These experiments revealed that MHC II is robustly expressed on subpopulations of myeloid progenitors and throughout the MK lineage in the bone marrow, including CD41+ MK biased HSCs, MK progenitors, and mature MKs.

**Fig. 1.**
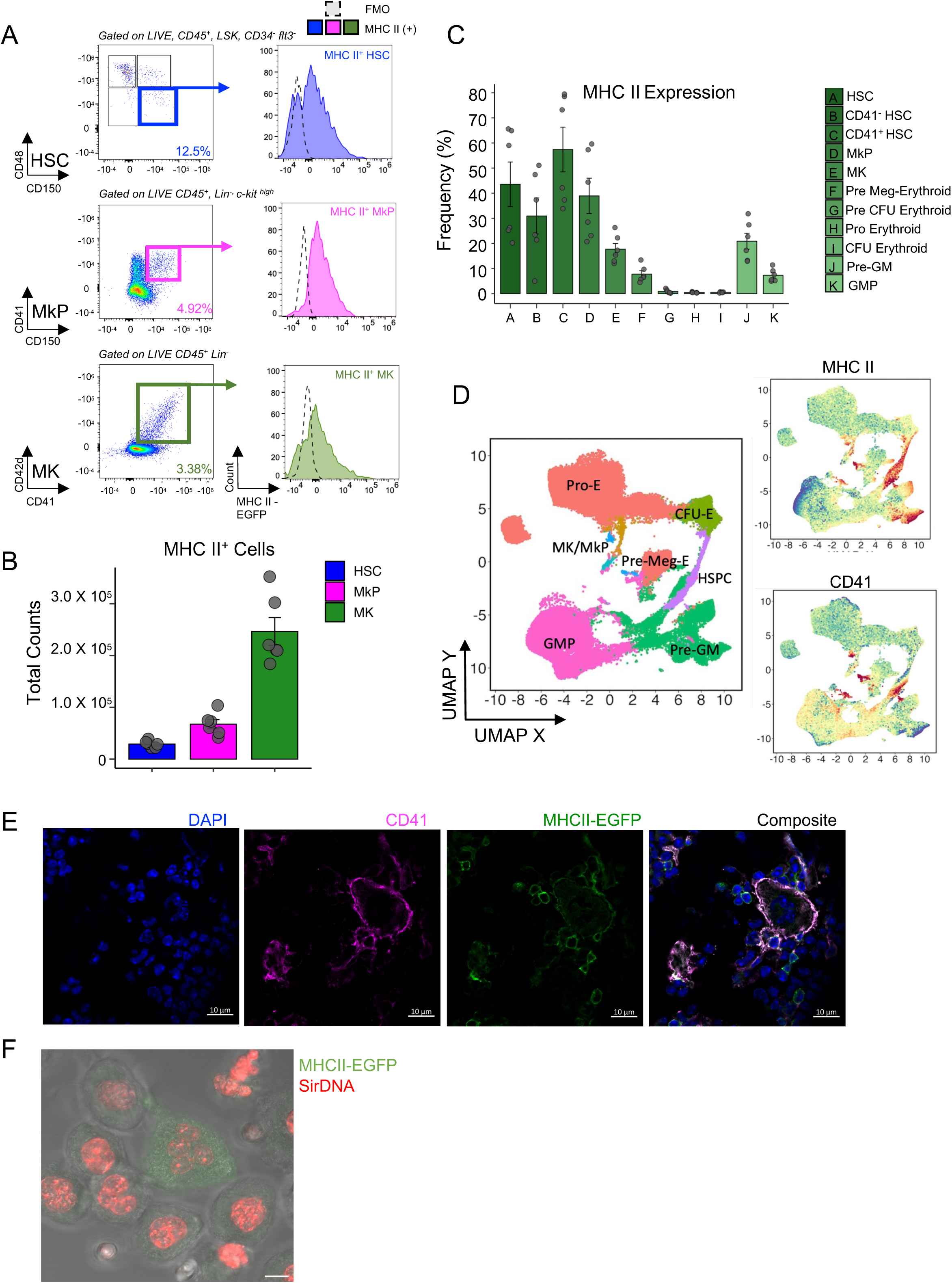
MHC II is expressed throughout the megakaryocyte lineage. (A) Representative plots and histogram of MHC II expression in HSCs, MkPs, and MKs. (B) Absolute counts of MHC II+ cells in murine hindlimb bones; *n* = 5 animals. (C) Frequencies of MHC II expression in bone marrow progenitors; *n* = 5 animals. (D) High dimensional flow cytometry UMAP projection of bone marrow progenitors (left); MHC II and CD41 expression (right). (E) Femoral cryosections stained with DAPI, CD41, MHC II; scale bar = 10 µm. (F) Representative images of native megakaryocytes isolated from MHC II-eGFP reporter mice with MHC II+ MKs visualized by endogenous eGFP expression (green) and nuclei stained with SiR-DNA (red); scale bar = 10 µm. Data are shown as mean ± SD; graphs represent data from at least 3 independent experiments.

### MHC II-expressing MKs activate CD4+ T cells ex vivo

We next aimed to determine the functional capacity of MK MHC II. MHC II molecules present peptides to CD4+ T helper cells, which in turn recognize the peptide-MHC II complex via their T-cell receptors (**TCRs**). For robust activation, however, a secondary signal between costimulatory molecules on both the CD4+ T cell and the APC must be provided. As such, we assessed if bone marrow MKs expressed key costimulatory molecules and found that they express CD80/CD86 (B7-1/B7-2), CD40, and CD83 at baseline (**Fig 2A**), indicating potential APC function. To ascertain if toll-like receptor (**TLR**) activation could influence the expression of the co-stimulatory molecules on MKs, we isolated native bone marrow MKs and treated them ex vivo with Polyinosinic-polycytidylic acid, (**Poly I:C**), a synthetic double-stranded RNA molecule that mimics viral RNA and activates TLR3 and Lipopolysaccharide (**LPS**), which activates TLR4. We observed that LPS led to upregulation of both MHC II and key co-stimulatory molecules on MKs upon stimulation (**Fig 2A**). Consequently, we next compared OVA uptake in MHC+ versus MHC-MKs and found a trend toward increased uptake within the MHC+ MK population (91.98 ± 8.45% vs. 76.05 ± 18.95%). This trend suggests that MHC II-expressing MKs may have greater capacity for OVA uptake; however, uptake was not restricted to the MHC II+ subset (**Fig S2A**). Given that key costimulatory molecules were upregulated upon TLR stimulation, and the high antigen-uptake capacity observed in MHC II+ cells, we next tested the ability of bone marrow MKs to process antigen through intracellular proteolysis using DQ-ovalbumin (**OVA-DQ**). MKs were treated with BODIPY-conjugated DQ-OVA, a self-quenched conjugate of OVA that exhibits bright green fluorescence only upon proteolytic cleavage, releasing the dye molecule from the OVA. After 90 minutes, DQ-OVA fluorescence was observed in MKs, indicating active intracellular proteolytic activity (**Fig S2B**). Thus, we next moved to functionally assay the ability of MKs to activate naive CD4+ T cells. We used OT-II mice engineered to express a TCR that recognizes specific residues of chicken OVA (OVA _323-339_) presented by MHC II. Bone marrow MKs were isolated, incubated with OVA peptide, and washed to remove unprocessed peptide before culture with naïve CD4+ OT II T cells (**Fig 2B**). We observed peptide-specific activation, as judged by CD25 upregulation, and proliferation of naïve OT-II T cells in conditions where both MKs and OVA were present (**Fig 2B-C**). Correspondingly, addition of an MHC II neutralizing antibody significantly abrogated T cell activation and proliferation (**Fig 2C**), demonstrating that T cell proliferation was an MHC II-dependent interaction. Live cell microscopy further substantiated the direct interactions between MKs and T cells (**Supplementary movie 1**). Given that T cell differentiation is also regulated by cytokine secretion, and that MKs are known to secrete TGF-β, we sought to determine how T cell interactions might be influencing this capacity. Indeed, we observed that after 5 days of coculture, MHC II-expressing MKs secreted more TGF-β, as quantified by intracellular LAP, in the presence of T cells (**Fig S2C**). As TGF-β drives induced Treg (**iTreg**) differentiation, we characterized the T cell compartment of the cultures. We observed that the presence of MKs increased the frequency of CD25+ Foxp3+ cells and that this was blunted with addition of a TGF-β neutralizing antibody (**Fig S2D**).

**Fig. 2.**
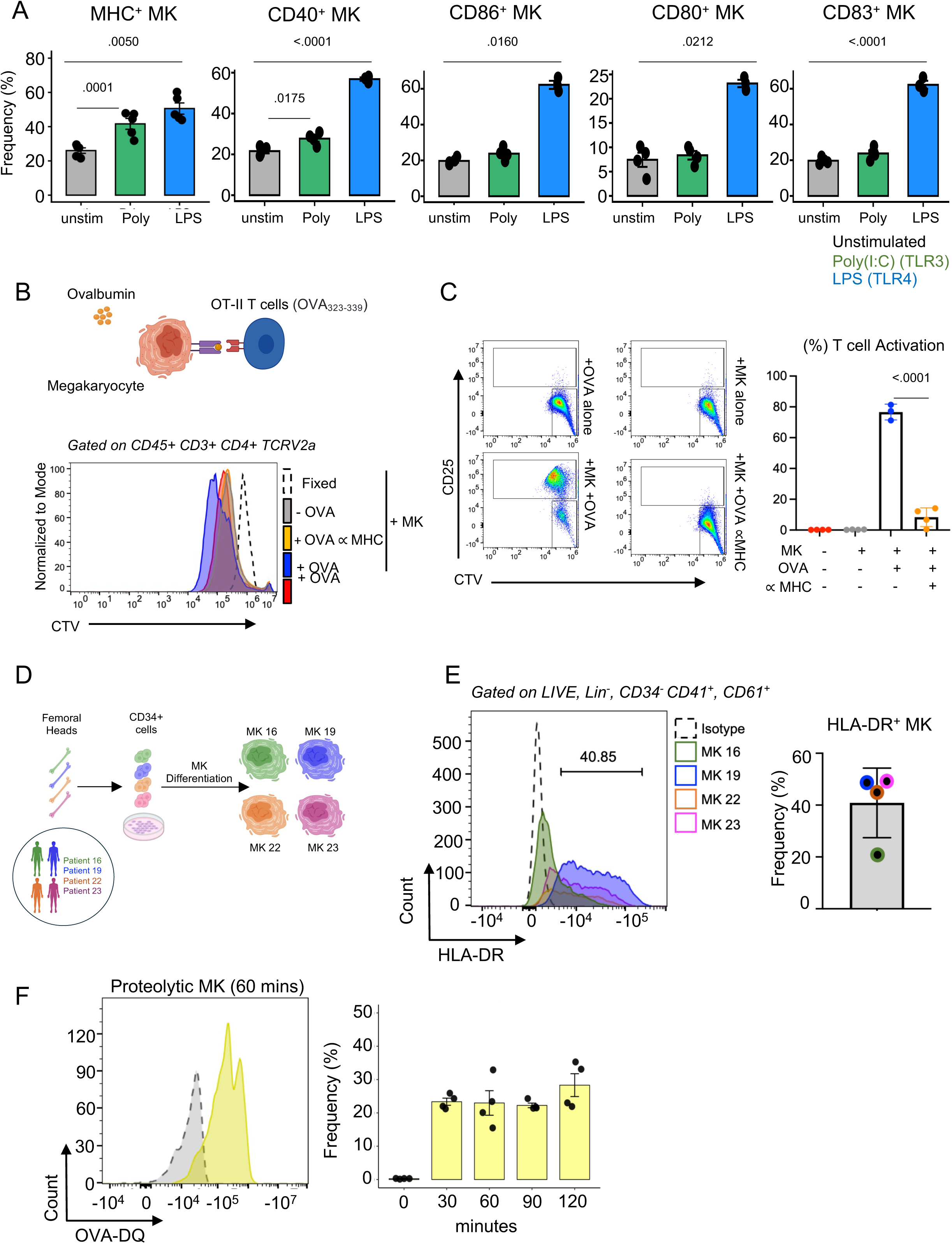
MHC II-positive megakaryocytes activate CD4+ T cells ex vivo. (A) Frequencies of CD40, CD86, CD80, and CD83 expressing MKs; *n* = 5 animals. (B) Schematic of murine bone marrow MK and naïve CD4+ T cell co-culture (top) and representative histogram of MK and T cell culture conditions in the presence or absence of OVA and anti-MHC antibody. (C) Representative plots of coculture conditions in the presence or absence of OVA and anti-MHC antibody (left); quantification of T cell activation (right). *n* = 4 mice/condition. (D) Schematic of ex vivo MK differentiation from human femoral head samples. (E) Representative histogram of HLA-DR+ human MKs (left) and frequency of expression (right). *n* = 4 donors. (F) Representative histogram of OVA -DQ+ fluorescent MKs (left) and frequency of OVA -DQ+ MKs at 30, 60, 90, and 120 mins (right); *n* = 4 donors. Data are shown as mean ± SD; graphs represent data from at least 3 independent experiments. Statistics performed with 1-way ANOVA with Tukey’s multiple comparisons test at 95% CI (A, C, F).

We next asked whether TLR stimulation of MKs altered the ability to influence other CD4+ T cell fates. Because TGF-β, particularly in combination with inflammatory cytokines, can also promote Th17 differentiation, we assessed IL-17A production; however, IL-17A was minimal across conditions, with no evidence of Th17 differentiation (**Fig S3A**). Following pre-stimulation with LPS or Poly I:C, we observed that MKs induced modest increases in CD4 IFN-γ and TNF-α T cells, with Poly I:C significantly increasing IFN-γ cells and LPS increasing TNF-α cells (**Fig S3A-B**). Specifically, when assessing Th1 or Th2 fates, we observed that stimulation with LPS or Poly I:C resulted in modest increases in Th1-associated responses, (T-bet+ IFN-γ+), and negligible Th2 differentiation (GATA3+ IL-4+)(**Fig S3A-B**).Together, these results demonstrate that bone marrow MKs activate naïve T cells in an MHC II-dependent manner, with MK/T cell interactions associated with increased TGF-β secretion and iTreg differentiation, while TLR stimulation of MKs elicits modest IFN-γ secretion.

To determine if our observations were applicable to human MKs, we isolated CD34+ cells from femoral head bone marrow samples obtained from healthy donors, which were differentiated into MKs (**Fig 2D**). We then characterized the cell surface expression of HLA-DR in ex vivo generated human MKs. The results mirrored our findings in the murine model; HLA-DR was expressed in approximately 40% of MKs (**Fig 2E**). To determine whether antigen-processing capacity was also a feature of human MKs, we incubated ex vivo-generated MKs from human bone marrow samples with DQ-OVA. We observed fluorescence resulting from OVA-DQ proteolysis, indicating that human MKs can process antigens, paralleling murine MKs (**Fig 2F**). To independently validate HLA-DR expression in primary human MKs, we analyzed circulating MKs within human PBMC samples by flow cytometry. These MKs showed HLA-DR expression of 52.6 ± 22.1%, comparable to that observed in bone marrow HSPC-derived MKs and supporting the relevance of this phenotype across human MK sources (**Fig. S3C**).

### MHC II is not present on platelets

We next further characterized the maturation state of MHC+ MKs and assessed if MHC II was transferred to platelets. As MKs mature, DNA replication occurs without cytokinesis in a process called endomitosis, leading to polyploidization. In turn, ploidy status correlates with maturation, and higher ploidy in combination with larger cytoplasmic content correlates with platelet production (13). Analysis of the ploidy status of bone marrow MKs revealed that MHC II expression spread across various ploidy states but predominated in lower ploidy MKs (**Fig 3A-B, S4A**). We and others have previously shown that MKs can transfer MHC I onto platelets, indicating that the machinery to transfer MHC components to platelets is present (18). Therefore, we examined if MHC Class II was present on platelets. First, we analyzed MK and platelet proteomes and compared the protein signature to acellular fractions: bone marrow fluid and peripheral blood plasma (**Fig S4B**). Principal component analysis (PCA) revealed that the proteomic profiles of bone marrow fluid and megakaryocytes (MKs) clustered most closely, indicating the greatest similarity between these compartments. In contrast, the largest differences were observed between platelets and bone marrow fluid. There were also significant differences between blood plasma and both MKs and bone marrow fluid **(Fig S4C**). To gain insight into proteins transferred from MKs to platelets, we first characterized proteins associated with either class I or class II antigen presentation function across platelets, MKs, blood plasma, and bone marrow fluid (**Fig 3C**). Visualization of antigen presentation proteins revealed proteomic overlap between MKs and platelets. These shared proteins likely represent factors synthesized in MKs and subsequently transferred to platelets during thrombopoiesis. In contrast, proteins detected exclusively in MKs or platelets were considered reflective of cell type-specific biological processes or differential protein turnover. Notably, components of the MHC class I pathway were among the shared proteins, suggesting that MHC class I molecules are transferred from MKs to platelets. Conversely, MHC class II-associated proteins were predominantly confined to MKs, consistent with limited inheritance of class II machinery by platelets (**Fig 3D**). These findings highlight the continuity between MK and platelet proteomes and suggest that platelets do not inherit MHC class II-associated proteins from precursor MKs.

**Fig. 3.**
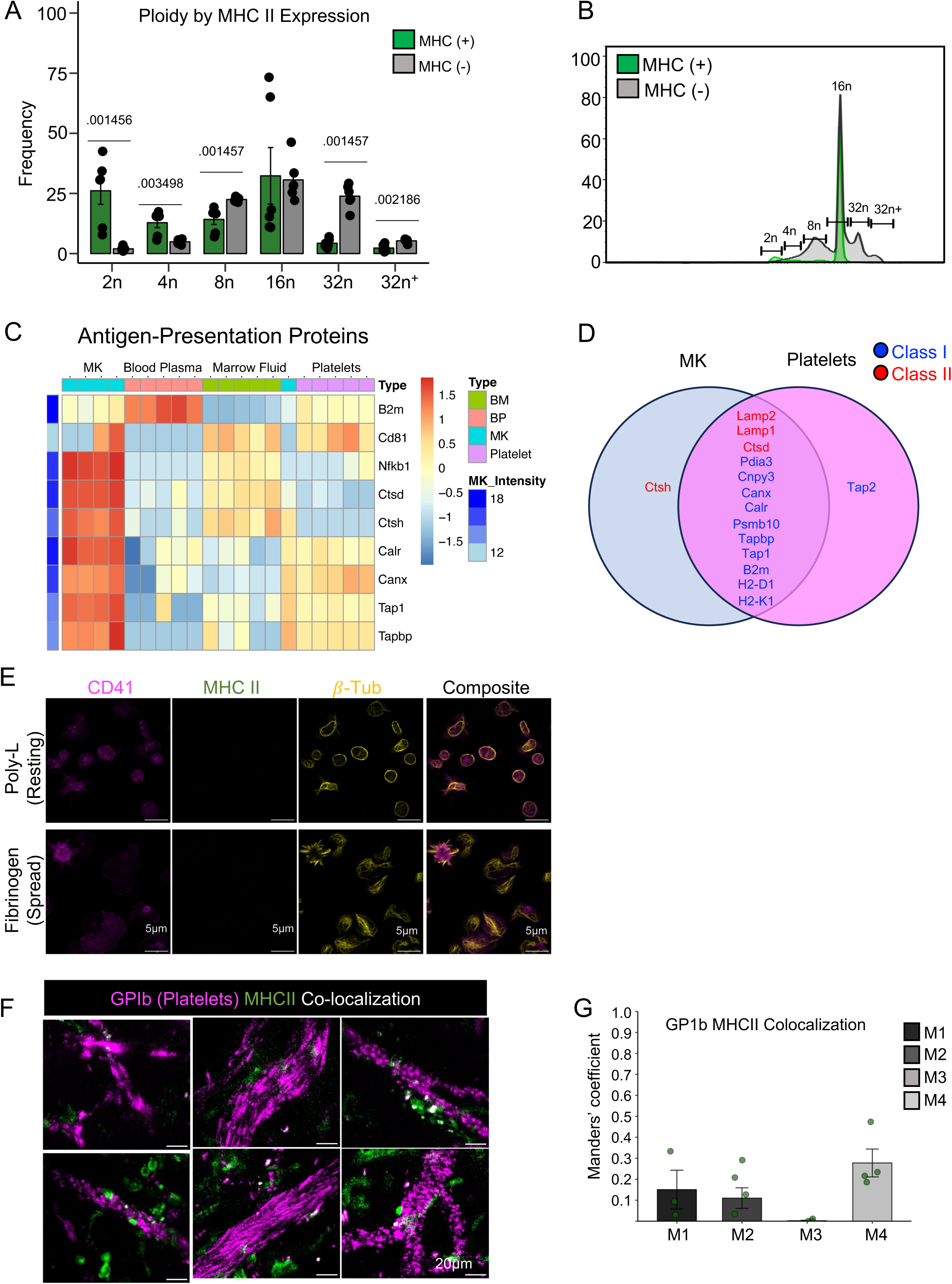
MHC II is present on megakaryocytes but not platelets. (A) Frequencies of DNA ploidy by MHC II expression; *n* = 10 animals. (B) Representative flow plot of MK ploidy distribution by MHC II expression. (C) Heatmap of proteins associated with antigen presentation in MKs, blood plasma, bone marrow fluid, and platelets; *n* = 4 animals. (D) Venn Diagram of MHC I and MHC II proteins shared between MKs and platelets: *n* = 4 animals. (E) Representative images of resting and spread platelets stained for CD41, MHC II, and β1-tubulin; scale bar = 5 µm. (F) Representative two-photon images of calvaria bone marrow from MHC II-eGFP reporter mice (green); platelets visualized by intravenous injection of fluorescently conjugated anti-GPIb antibody (magenta). Areas of colocalization between MHC II+ cells and platelets are shown in white; scale bar = 20 µm. (G) Quantification of platelet and MHC II colocalization. N = 4 mice, with 3–5 fields of view analyzed per mouse. Data are shown as mean ± SD (A); mean ± SEM (G); graphs represent data from at least 3 independent experiments (A-B, F-G). Data from 1 independent experiment with 4 biological replicates (C-D). Statistics performed with unpaired 2-tailed Student *t* test.

To expand on this, we performed immunostaining of MHC II on resting and activated platelets. We detected no MHC II signal in either condition (**Fig 3E**). To validate this in vivo, we performed two-photon intravital microscopy (2P-IVM) on the calvaria of MHC II-EGFP knock-in mice. To mimic immune activation, mice were treated with LPS, which simulates immune activation via TLR4. However, even upon TLR stimulation, no MHC II was detected in platelets, as assayed by colocalization of GFP and GP1B. (**Fig 3F-G**, **Supplementary Movies 2-3**). Together, these data suggest that MHC II-expressing MKs are either not platelet-producing, or do not package this molecule into platelets, even when MHC II is upregulated.

### MHC II+ vs MHC II-MKs have differential gene expression

To further define the MHC II+ MK population, we performed bulk RNA sequencing (**RNA-seq**) of MHC II-positive and -negative MKs. We observed various DE genes between the two subsets while pathway analysis showed an enrichment of functions related to phagocytosis, interferon responses, and cell-adhesion in MHC II-positive MKs. Conversely, MHC II-negative MKs were enriched in functions related to migration, chemotaxis, and coagulation (**Fig 4A**, **S5A-B**, **Supplementary Table 1**).

**Fig. 4.**
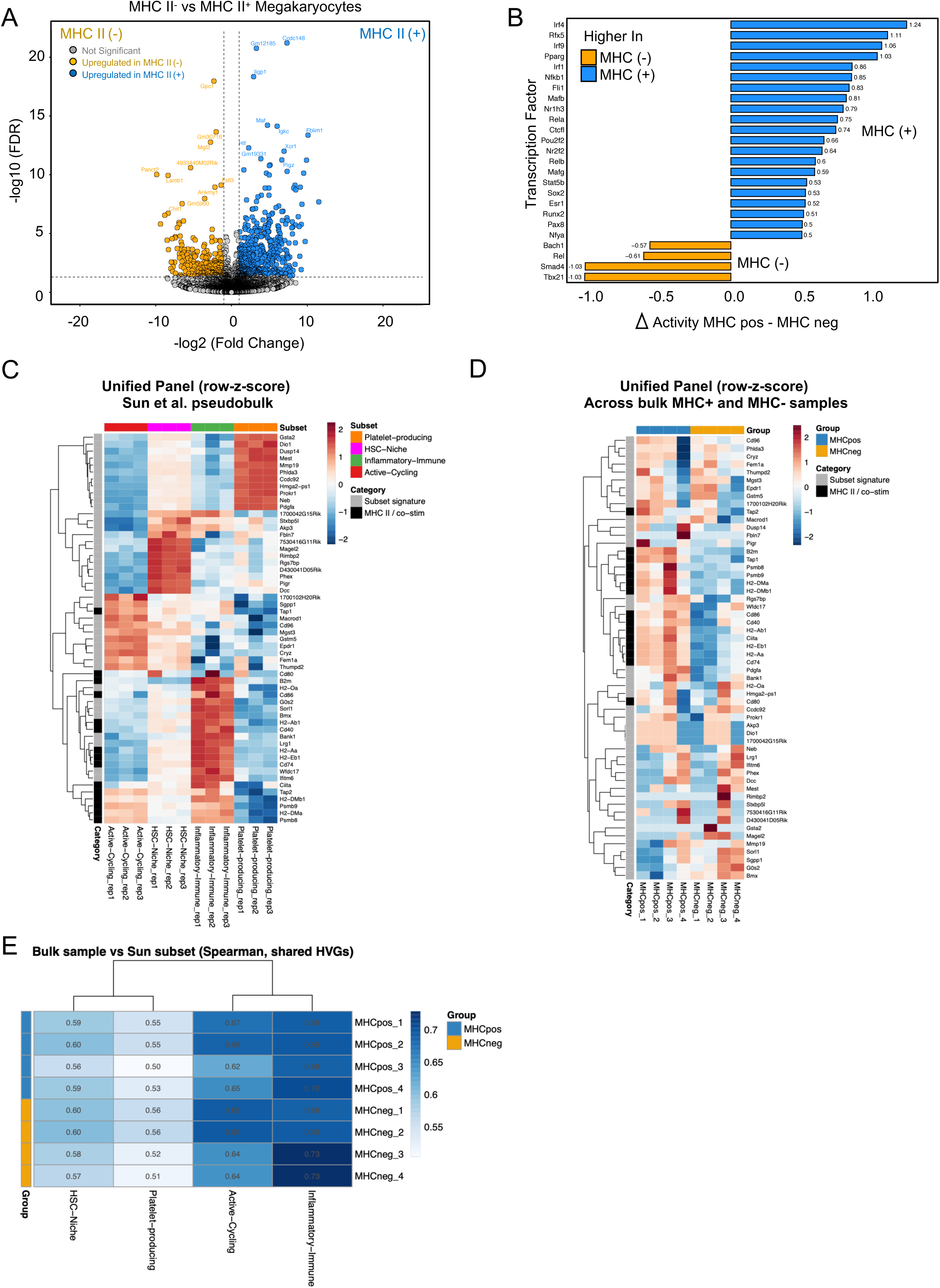
MHC II-positive megakaryocytes have a unique transcriptome enriched in interferon regulatory factors and immune-related transcriptional programs. (A) Volcano plot of differentially expressed genes between MHC II- and MHC+ MKs. (B) Bar plot of DE genes integrated with (TF) regulons from the DoRothEA database showing the top 35 TFs ranked by differential activity (Δ activity = MHC+ – MHC-). Bars are colored according to the group in which TF activity was higher (blue, MHC+; orange, MHC-), with values indicating the magnitude of difference in inferred activity. (C) Integration of bulk MHC II+/- MK transcriptomes with the Sun et al. (2021) single-cell MK atlas (four subsets, marker-annotated; cells per subset: Active-Cycling 343, HSC-Niche 271, Inflammatory/Immune 143, Platelet-producing 38). Row z-scored expression of a 56-gene unified panel (subset-defining signatures plus MHC class II and co-stimulatory module) across pseudobulk profiles of the four reference subsets (three pseudo-replicates each). (D) The same panel across the bulk MHC II+ and MHC II-samples (n = 4 per group). (E) Spearman correlation of each bulk replicate to each reference-subset pseudobulk over 2,000 shared highly variable genes; cell values are correlation coefficients.

To identify the transcriptional regulators driving the differential gene expression between MHC II-positive and MHC II-negative MKs, we employed VIPER analysis in combination with the DoRothEA regulon database. Analysis of the inferred transcription factor (**TF**) activities revealed distinct sets of TFs with elevated activity in each population. In MHC II+ MKs, the top upregulated TFs included Irf4, Rfx5, Irf9, Pparg, and Irf1 (**Fig 4B**, **S5C**). Predictably, Rfx5 serves as a regulator of MHC class II genes, directly driving the expression of antigen presentation machinery; Irfs suggest involvement in immune signaling and antiviral responses, while Pparg indicates involvement in lipid metabolism and potential anti-inflammatory pathways. Overall, this network indicates a strong involvement of interferon regulatory factors and immune-related transcriptional programs.

Conversely, in MHC II-cells, TFs such as Smad4, Tbx21, Bach1, and Rel exhibited the highest activity, highlighting transcriptional programs potentially involved in TGF-β signaling, oxidative stress response, and NF-κB signaling (**Fig 4B**, **S5C**). This analysis suggests that different transcription factors likely orchestrate the transcriptional differences between MHC II-positive and -negative MKs and reveals that the activity of specific interferon-regulated and immune-modulatory TFs is a defining feature of MHC II+ cells.

Previous work by Sun et al. defined four transcriptional subsets of bone marrow MKs: platelet-generating, HSC-niche, inflammatory/immune, and actively cycling. To place our MHC II+ and MHC II-transcriptomes in this framework, we generated pseudobulk profiles for each subset from the Sun et al. atlas and compared them with our bulk data across a 56-gene unified panel combining the four subset signatures with MHC class II and co-stimulatory genes (**Fig 4C**). MHC class II and co-stimulatory genes (H2-Ab1, H2-Aa, Cd74, Cd80, Cd86, Cd40) were expressed most highly within the inflammatory/immune subset but were also detectable across other subsets (**Fig 4D**). Quantitative mapping by rank correlation over shared highly variable genes assigned all four MHC II+ replicates to the inflammatory/immune subset (rho 0.68 to 0.70), whereas MHC II-replicates were distributed between the inflammatory/immune and actively-cycling programs (**Fig 4E**). Both populations correlated least with the platelet-producing and HSC-niche subsets (**Fig 4C-D**). The specificity of this mapping was confirmed by a pseudobulk control in which all reference subsets were correctly re-identified (**Fig S5D**). This was further validated by single cell scoring of our MHC signatures across the Sun et al. subsets (**Fig S5E**) and mapped into original dataset (**Fig S5F**). We therefore interpret MHC class II expression not as the property of a discrete MK subset but as an immune-active transcriptional state centered on the inflammatory/immune program, consistent with antigen-presentation potential being enriched within, rather than exclusive to, a single MK subset.

### MK MHC II is peptide-occupied in vivo

To substantiate that the MHC II on MKs is occupied in vivo and characterize the MHC II-associated peptidome of bone marrow MKs, we performed immunopeptidomics of in situ MKs. We employed immuno-capture of MHC II complexes followed by mass spectrometric analysis of eluted peptides, which enabled the identification of MHC II-bound peptides from bone marrow MKs (**Fig 5A**). High-resolution LC-MS/MS of sorted bone marrow MKs identified 290 peptides corresponding to 75 proteins. After filtering for peptide length (≥9 residues), removal of contaminants, and exclusion of MHC-II-derived sequences, the dataset consisted of 78 peptides and 43 parental proteins (**Supplementary Table 2**). Peptide length distribution showed that over 90% were 9–26 amino acids, with a mode of 15 residues, consistent with canonical MHC class II ligands (**Fig S6A**). GibbsCluster v2.0 was used for unsupervised motif analysis of the peptide dataset. The GibbsCluster output for the peptide dataset showed good agreement with the H2-Ab1 motif described in the database (**Fig S6B**), supporting the assignment of these peptides as H2-Ab1-associated ligands. To predict the theoretical binding affinity of peptides to murine H2-Ab1 molecules, we utilized the NetMHCIIpan v4.1 tool. Using a 1% rank threshold for strong binders and a 5% rank threshold for weak binders, 51.3% of peptides were classified as strong binders and 5.1% as weak binders (**Fig 5B**). Mapping peptides to their source proteins revealed that a small number of proteins dominated the immunopeptidome. The most abundant was Ferritin light chain 1 (Ftl1), followed by Vimentin (Vim), Adenylyl cyclase-associated protein 1 (Cap1), and Prolow-density lipoprotein receptor-related protein 1 (Lrp1) (**Fig 5C-D**). Nested sets, groups of peptides sharing a common 9-mer binding core but varying in C- and N-terminal lengths, are common among MHC II peptides. We categorized these sequences into clusters which provided a visual approximation of the peptide families present in the dataset. Thirteen clusters were identified, with a notable group being a set of nine peptides derived from the ferritin light chain 1 protein (**Fig 5D**). Classification into protein categories revealed an enrichment of metabolism (52.9%) and cytoskeleton (22.1%) associated proteins (**Fig 5E**). To explore the functional enrichment of the 43 parental proteins from MHC II-isolated peptides, a GO enrichment analysis was performed using annotations from the PANTHER database. The proteins were categorized based on their molecular functions, biological roles, and protein classes. The molecular function analysis highlighted a significant enrichment in binding activities. In terms of biological processes, the main enrichments were in cellular processes and biological regulation (**Fig S6C**). Lastly, the analysis identified protein-binding activity modulators as the most prevalent protein classes within the group (**Fig S6C**).

**Fig. 5.**
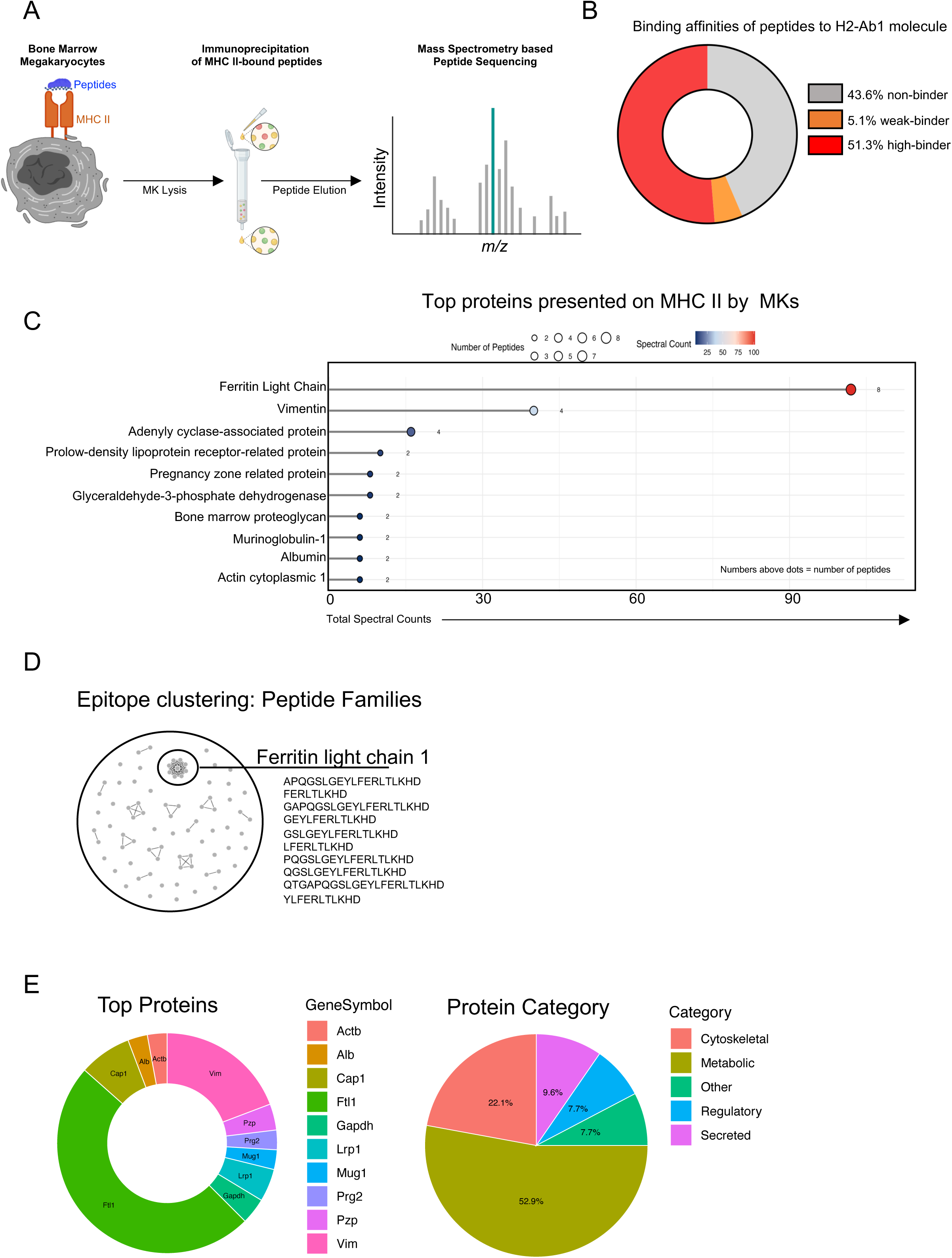
Immunopeptidomics reveals MHC II on native bone marrow megakaryocytes is occupied with peptide. (A) Schematic of bone marrow MK MHC II immunopeptidomics. (B) Donut plot displaying binding affinities of MK peptides to the H2-Ab1 molecule. The cumulative percentage of high-binder (HB), weak-binder (WB) and non-binder (NB) isolated peptides is shown. (C) Lollipop plot of spectral counts displaying number of peptides in respective protein families. (D) Epitope cluster analysis of MK peptides grouped into clusters based on sequence identity, by applying a threshold of 70%. (E) Donut plot displaying top proteins identified (left) and assigned to protein category (right).

MHC II molecules primarily present peptides derived from exogenous proteins. Proteins are internalized by several pathways, including phagocytosis, macropinocytosis, and endocytosis, and peptides are generated by lysosomal proteolysis (19, 20). Thus, endo-lysosomal degradation pathway includes proteins from cell membrane, extracellular region, Golgi apparatus, endoplasmic reticulum, and endosome/lysosome, while cytosolic pathway of degradation includes cytosolic, mitochondrial, and nuclear proteins. Of the parental proteins, 62.8% were annotated to endo-lysosomal compartments or trafficking pathways, whereas the remainder were associated with cytosolic, mitochondrial, or nuclear compartments (**Fig. S6D**). These annotations are consistent with contributions from exogenous and endogenous sources, although peptide origin cannot be definitively assigned without tracer-based experiments

### MK-specific loss of MHC II alters stem and progenitor cells following TLR stimulation

To better understand the functional differences between MHC II-expressing and non-expressing MKs and discern the relevance of the MHC II-expressing MKs, we generated a murine model with targeted deletion of MHC II in the MK lineage*, (Pf4-MHC*^Δ/Δ^ mice; MHC^KO^ mice). When analyzed by flow cytometry, we observed deletion of MHC II in mature MKs, partial deletion in MkPs and minimal loss of MHC II in HSCs (**Fig S7A**). *Pf4-MHC*^Δ/Δ^ mice had no change in MK numbers as quantified by immunofluorescence of femoral cryosections and no change in MK size (**Fig 6A**). Notably, the genetic deletion of MHC II functionally validated the essential role of MK-derived MHC II in antigen presentation, as MKs from MHC^KO^ mice lost the ability to present antigen and activate naïve CD4+ T cells (**Fig S7B**). We also observed that in steady state conditions MHC II^KO^ MKs had decreased TGF-β1 production as quantified by LAP staining of femoral cryosections (**Fig 6B**) aligning with ex vivo data showing MHC II-expressing MKs produced more TGF-β after co-culture of MKs with T cells (**Fig S2C**).

**Fig. 6.**
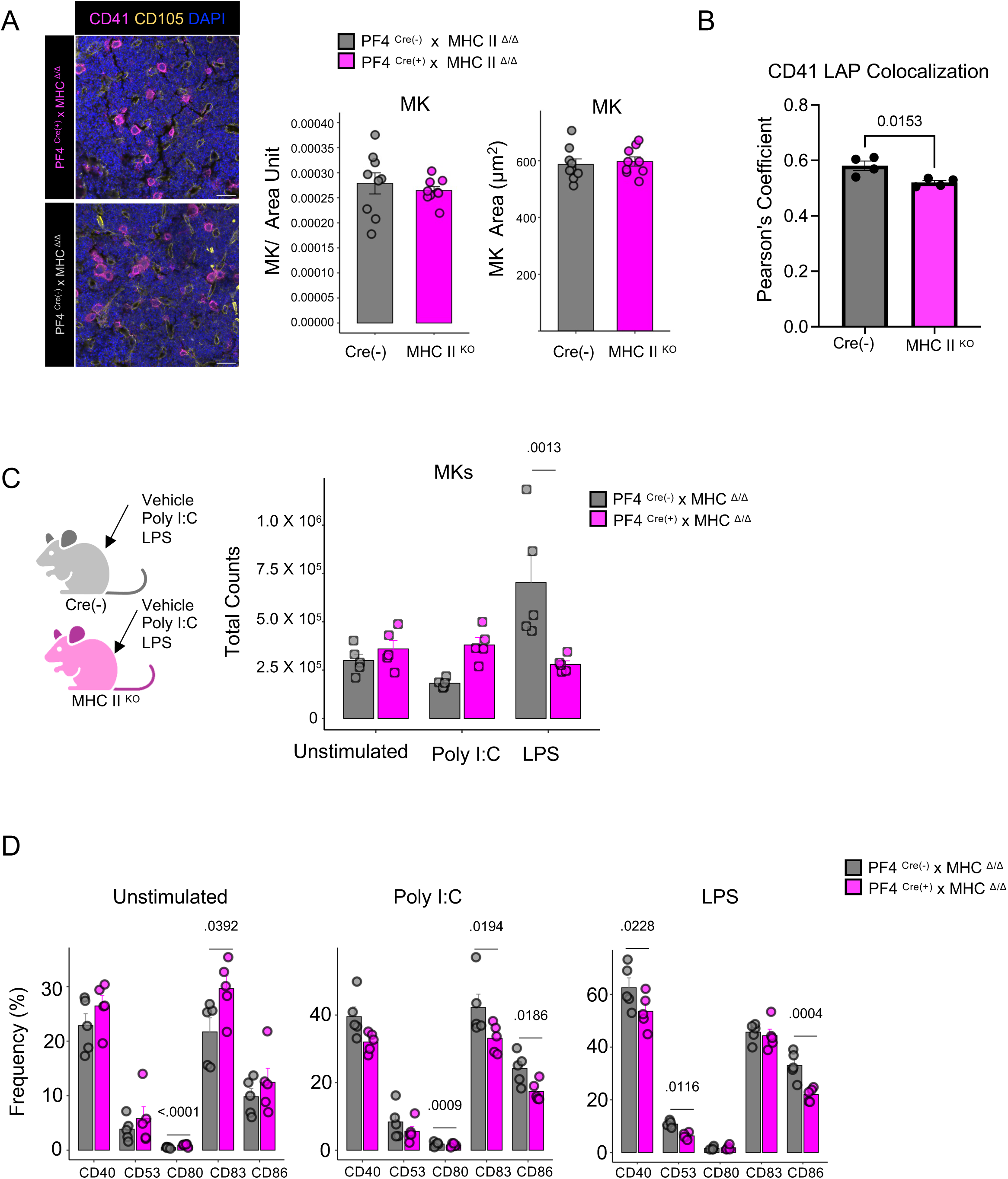
Megakaryocyte-specific loss of MHC II alters megakaryocyte TLR sensitivity. (A) Representative images (left) and quantification of MK numbers and MK area (right) in MHC^KO^ and Cre (^-^) mice; *n* =10 mice/ genotype. (B) Colocalization analysis of LAP and CD41 on femoral cryosections of MHC^KO^ and Cre (^-^) mice; *n* =4 mice/ genotype. (C) Total MK counts in murine hindlimb bones 24 hours after LPS or Poly (I:C), or vehicle administration; n= 5 mice/ genotype. (D) Frequency of CD40, CD53, CD80, CD83, CD86 (+) MKs 24 hours after LPS, Poly (I:C), or vehicle administration; *n* = 5 mice/ genotype. Data are shown as mean ± SEM; graphs represent data from at least 3 independent experiments. Statistics performed with unpaired 2-tailed Student *t* test (A-B) and 2-way ANOVA with Tukey’s multiple comparisons test at 95.00% CI (C-D).

Next, we administered in vivo LPS and poly(I:C) to determine if MHC II expression affected MK responses to inflammation. Deletion of MHC II on MKs differentially altered MK expansion and costimulatory molecule expression. Specifically, following LPS administration, MK numbers were lower in *Pf4-MHC*^Δ/Δ^ mice. Conversely, MKs expanded in response to poly(I:C) in *Pf4-MHC*^Δ/Δ^ mice (**Fig 6C**). MKs lacking MHC II also had altered expression of costimulatory molecules both at baseline and after stimulation with LPS and poly(I:C) (**Fig 6D**). These findings suggest that MHC II expression on MKs modulates their inflammatory response in a stimulus-dependent manner, influencing both MK expansion and the expression of costimulatory molecules.

We next examined if deletion of MK MHC II had any effects on stem and progenitor populations, as MKs are also active regulators of the bone marrow niche, and the role of MHC II-expressing MKs on HSPCs is largely unknown. We uncovered a minor decrease in HSCs and ST-HSCs in *Pf4-MHC*^Δ/Δ^ mice but did not detect a change in HSC proliferation (**Fig S7D**). In addition, loss of MHC II in MKs resulted in additional HSPC population alterations in response to TLR stimulation. In *Pf4-MHC*^Δ/Δ^ mice treated with LPS, the HSC pool was resistant to depletion, while there was a significant increase in MkP, MPP2, and MPP4 populations. In response to Poly I:C, there was also a slight increase in MPP2 cells (**Fig S7E**). These data suggest that MHC II-expressing MKs may help shape inflammatory remodeling of the bone marrow stem and progenitor compartment.

## DISCUSSION

In this study, we characterized the expression and functionality of MHC II in bone marrow MKs. We demonstrate that a subset of MKs express MHC II and co-stimulatory molecules, are capable of antigen uptake and processing, and can activate naïve CD4+ T cells in an MHC II-dependent manner ex vivo. Notably, MHC II was not detected in platelets, even when platelets were activated. This contrasts with MHC Class I, which is transferred from MKs to platelets and modulates CD8+ T cell responses (18). Using an MHC II-EGFP knock-in mouse and a murine model with targeted deletion of MHC II in the MK lineage, our work expands the understanding of MKs beyond their platelet-producing functions and positions them as immune regulators in the bone marrow.

Our data revealed that bone marrow MKs express not only MHC II but also co-stimulatory molecules such as CD40, CD83, CD80, and CD86. This aligns with transcriptional analyses identifying an “immune MK” subset enriched in antigen presentation and TLR signaling genes (2, 8). Our data indicate that MHC II+ MKs are transcriptionally centered on the inflammatory/immune program described by Sun et al., whereas MHC II-MKs span a broader immune and cycling profile, indicating that immune competence can be engaged across overlapping MK populations. Treatment with the TLR agonist LPS enhanced MHC II and co-stimulatory molecule expression on MKs, indicating a functional response. We also established that bone marrow MKs activate CD4+ T cells ex vivo in a peptide-specific manner. Crucially, T cell proliferation was abrogated by MHC II blockade, confirming the dependence on this receptor. The relevance of these findings extends to human MKs, where similar expression of the MHC II molecule HLA-DR was observed in approximately 40% of human MKs differentiated from healthy donors. Additionally, the capacity for antigen processing was confirmed using a self-quenched DQ-ovalbumin assay, demonstrating that human MKs can internalize and process exogenous antigen. Together, these data establish that MKs possess the hallmarks of APCs and may modulate immune responses in the bone marrow.

To characterize the peptide repertoire presented by MK MHC II under homeostatic conditions, we performed immunopeptidomics. The predominance of Ftl1 and Vim within the MK MHC II immunopeptidome suggests that iron homeostasis and cytoskeletal integrity may be among the cellular processes visible to immune surveillance in the bone marrow. Additional enriched proteins, such as Cap1 and Lrp1, point to potential contributions from vesicular trafficking and lipid metabolism. Overall, these findings indicate that MHC II molecules in MKs capture a mixture of structural, metabolic, and regulatory proteins. However, the prominence of these peptides could also result from protein abundance or intrinsic peptide-MHC affinity. Moreover, the detection of peptides from these proteins does not establish that the proteins themselves participate in antigen presentation pathways. Further work is necessary to determine how specific peptides influence CD4+ T cell behavior within the marrow niche.

The bone marrow is increasingly recognized as an immunologically active niche. Our findings, together with work by Hernández-Malmierca et al. (8) suggest that antigen presentation by atypical APCs within the bone marrow is a mechanism for safeguarding hematopoiesis via direct interactions with CD4+ T cells. Our data extend this, suggesting that MHC II+ MKs can additionally engage CD4+ T cells. Further, we found that *Pf4-MHC*^Δ/Δ^ mice had a decrease in bone marrow TGF-β. Future studies examining Treg function and activity, together with broader profiling of T cell populations in the bone marrow, will be important to further define the immunoregulatory consequences of reduced MK-derived TGF-β or other cytokines.

We also found that MK-specific deletion of MHC II altered MK responsiveness to TLR agonists. MHC^KO^ MKs had an increased baseline expression of several costimulatory molecules, with the most significant differences in CD80 and CD86. However, upon stimulation with LPS and poly(I:C), MKs lacking MHC II had decreased upregulation of CD80, CD83, and CD86. This reduced induction of costimulatory molecules in MHC II-deficient MKs after TLR stimulation is likely driven by broader, cell-extrinsic changes including the absence of MHC II-dependent feedback from CD4+ T cells or shifts in the surrounding immune landscape, both of which normally amplify TLR-driven costimulatory programs.

This study also has limitations. Our results establish that MKs can activate naive CD4+ T cells to differentiate into Tregs ex vivo via MHC II and TGF-β, and that MK MHC II is peptide-occupied in vivo. However, additional work is necessary to determine if MKs mediate antigen-specific CD4+ T cell priming in vivo. Moreover, while our proteomic analysis and imaging did not detect MHC II on platelets, these approaches may not exclude extremely rare or transient expression on newly released platelet subsets and/or in specific disease conditions. In our immunopeptidomics analysis, peptide length distribution showed that over 90% were 9-26 amino acids, with a mode of 15 residues (**Fig S6A**), and motif analysis agreed with H2-Ab1 (**Fig S6B**), supporting bona fide class II ligands. While GO-based endo-lysosomal annotations (**Fig S6D**) align with exogenous processing routes, these compartments also handle endogenous membrane and organelle turnover; thus, source attribution remains inferential without in vivo tracers. Another limitation of this study is that the functional T cell co-culture assays were performed using ex vivo TPO-differentiated MKs rather than freshly isolated bone marrow MKs. This approach enabled the generation of highly purified, viable MKs in sufficient numbers for mechanistic studies while minimizing contamination by other antigen-presenting cells. Future studies using freshly isolated bone marrow MKs will be valuable to further validate these findings. There are also limitations to our *Pf4-MHC*^Δ/Δ^ mouse model. Given the minimal HSC deletion observed (**Fig S7A**) and the expression of *PF4* in the HSC compartment we cannot fully exclude direct HSC-intrinsic contributions to HSPC phenotypes and TLR responses in *Pf4-MHC*^Δ/Δ^ mice or the fact that deletion in a small number of HSC results in partial deletion across all other differentiated lineages stemming from this pool (21). Finally, we also note that the sorting and enrichment procedure may reduce the recovery of larger, highly polyploid megakaryocytes, and therefore the highest-ploidy MKs (both MHC+ and MHC-) may be underrepresented in our transcriptional analysis.

In summary, our work suggests that MHC II+ MKs may influence adaptive immunity in the bone marrow. Together with recent findings (9, 22) demonstrating antigen presentation as a safeguard for HSC integrity, our results expand the concept of “immune-active” MKs and hematopoietic cells, suggesting potential avenues for therapeutic targeting in immune-related conditions.

## MATERIALS AND METHODS

### Study design

This study was designed to investigate the function and phenotype of MHC II-expressing megakaryocytes (MKs) in the bone marrow. Methods include immunopeptidomics, bulk RNA-seq, flow cytometry, immunofluorescence microscopy, confocal imaging, live cell imaging, and ex vivo culture systems. Details of biological replicates, experimental replicates, and statistical analyses are provided in the corresponding figure legends. All details of experimental parameters are provided in Materials and Methods or the Supplementary Materials sections. For image analyses, the study design was blinded to investigators.

### Murine strains

C57BL6 mice (stock no. 000664), OT-II mice (stock no. 004194), Pf4-Cre mice (stock no. 008535), *MHCII ^fl/fl^*mice (stock no. 037709), and *Foxp3*^EGFP^ mice (stock no 006772) were obtained from The Jackson Laboratory. *I-A^b^*β-eGFP mice were described previously and were bred in the Harvard Medical School animal facility (15).

### Immunopeptidomics: Megakaryocyte sort and lysis

Bone marrow MKs were sorted (Live Lineage^−^ CD45^+^ SYTO-40^+^, CD41^+^CD42d^+^) from 10 to 12-week-old B6 mice into PBS. Pellets were snap frozen. Samples were pooled and lysed/solubilized as follows: 500 µL of 50 mM Tris-HCl, 150 mM NaCl (pH 8.0), 1x complete protease inhibitor cocktail, and 1% n-Dodecyl-β-D-maltoside (DDM) were added to sample 1, and 500 µL of 50 mM Tris-HCl, 150 mM NaCl (pH 8.0), 1x complete protease inhibitor cocktail, and 1% NP-40 were added to sample 2. Four cycles of 5-second sonication/5-second break were carried out on ice. Samples were incubated for 30 min in rotation at 4 °C, followed by centrifugation at 20,000 x g for 1 h at 4 °C. Supernatants were collected and one aliquot was taken for western blot (WB) analysis (2%), and one for protein quantification by BCA. WB conditions were as follows: Primary antibody; Hamster/mouse anti-mouse MHC Class II (β chain), clone KL277 (1/500; o/n incubation) and Secondary antibody; Fluorescent goat anti-mouse IgG (1/10,000; 1h incubation).

### Immunopeptidomics: Immunoprecipitation (IP)

The supernatant containing solubilized membranes (SB) was incubated overnight (o/n) at 4 °C, with an anti-mouse MHC Class II (I-A/I-E) antibody (Ab), clone M5/114, coupled to CNBr-activated sepharose beads: 20 µL of a 0.5x preparation of Ab-beads coupled at 2 mg Ab per 1 mL of hydrated beads were added. After o/n incubation, sample was loaded into a Bio-Spin Chromatography Column (Bio-Rad). Non-retained (NR) fraction was collected and sequential washes were performed on the retained fraction: 3x 50 mM Tris-HCl, 150 mM NaCl (pH 8.0), 0.5% DDM; 3x 50mM Tris-HCl, 150 mM NaCl (pH 8.0); 1x 50 mM Tris-HCl, 0.5M NaCl (pH 8.0); 3x 50 mM Tris-HCl, 150 mM NaCl (pH 8.0); and 2x 20 mM Tris-HCl (pH 8.0). Peptides were eluted with 0.25% trifluoroacetic acid (TFA) and stored at -80 °C. Aliquots of 5% NR and 10% of eluates (EL) were collected for WB analysis to assess the yield of IP.

### Immunopeptidomics: Mass spectrometry (MS) analysis

Sample was evaporated to dryness, reconstituted in 2.5% TFA, and loaded into ZiptipsC18 pipette tips for desalting. Sample was then fractionated by strong cation-exchange chromatography using GELoader epTIPS loaded in house with polysulfoethyl-A. Elution with NH_4_Cl at 25 and 75 mM in 30% acetonitrile (I) / 0.1% formic acid (FA) was applied. The fractions were evaporated to dryness and reconstituted in 5% methanol / 0.5% TFA for MS analysis. The MS system used was a 480 Exploris mass spectrometer (ThermoFisher) coupled to ACQUITY UPLC M-Class chromatographic system (Waters). The separation was conducted at 500 nL/min, gradient 0-30% ACN (0.1% FA) in 60 min. The spectrometric analysis was performed in a data dependent mode, HCD fragmentation, Top speed method. The range acquired was 400-1,600 m/z. LC-MS/MS spectra were searched using SEQUEST (Proteome Discoverer v3.0, ThermoFisher) with the following parameters: peptide mass tolerance 10 ppm, fragment tolerance 0.02 Da, and dynamic modification of methionine oxidation (+15.995 Da). The database used for searching was created by combination of *Mus musculus* (Uniprot), and common proteomics contaminants database (MaxQuant). After filtering results by high peptide confidence (1% FDR), 290 peptides and 75 proteins were obtained from database search.

### Proliferation Assay via EdU

A total of 1 mg/mouse of EdU in 200 μL PBS was administered intraperitoneally (i.p.) 48 hours before being euthanized for analysis. Cells were analyzed 48 hours after injection, and EdU incorporation was assessed according to the manufacturer’s protocol.

### In Vivo TLR Stimulation

A total of 10 mg/kg of poly (I:C) or 250 ug/kg of LPS in 200 uL of PBS was administered intraperitoneally (i.p.) to mice 24 hours before being euthanized for analysis. Cell populations were analyzed 24 hours after injection.

### Ex Vivo MK TLR Stimulation

Cultured megakaryocytes were enriched by size exclusion and stimulated with either LPS (10 ng/mL), poly(I) (1 µg/mL), or vehicle for 4 hours. Following stimulation, cells were washed with media containing 10% serum to remove residual LPS or poly(I) prior to addition to T-cell cultures.

### Flow cytometry and cell sorting

Cell surface staining was performed following Fc blocking (TruStain FcX™, anti-mouse CD16/32) by incubating single cell suspensions. Cells were then stained with surface markers. A viability dye (Life Technologies, Aqua, dilution 1:1000) was applied to exclude dead cells and SYTO™ 40 Blue (S1135, ThermoFisher Scientific) was used to indicate viable cells. MKs were sorted using a 130 um nozzle and BD FACSymphony™ S6 Cell Sorter. For immunophenotyping, data was acquired and unmixed using SpectroFlo^®^ v2.2.0.3 software (Cytek Biosciences, Fremont, California, USA). FlowJo (Tree Star Inc.) was used for data analysis. High-dimensional analysis was conducted using *Spectre* toolkit in R (v.4.5.1).

### Megakaryocyte isolation by size exclusion

Bone marrow cells hind limb bones (femora, tibiae, and iliac crests) were isolated by centrifugation for 40 seconds at 2000 x g and primary MKs were isolated as follows. Red blood cells were lysed for 5 mins in ACK buffer (A1049201, Gibco) and cells were passed through a 100 µm cell strainer. The cell suspension was consecutively filtered using 20 and 15 µm cell strainers (43-50020-03 and 43-50015-03, pluriSelect) and MKs were retrieved by inverting strainers, as published (23).

### Murine megakaryocyte differentiation

MKs were differentiated from murine lineage depleted cells harvested from femurs and tibias of 8–12-week-old B6 mice. Progenitor cells from the bone marrow cell suspension were enriched by depleting mature, lineage^+^ cells expressing CD3ε (clone 145-2C11), CD19 (clone 1D3), B220 (clone RA3-6B2), NK1.1 (clone PK136), CD11b (clone M1/70), CD11c (clone N418), Ly6G/C (clone RB6-8C5), and/or Ter119 (clone TER-119) followed by magnetic bead isolation using DynabeadsTM (11415D, ThermoScientific). Enriched progenitor cells were supplemented with 50 ng mL^-1^ TPO (DMEM, 10% FBS, 2 mM glutamine, 100 IU/mL penicillin/streptomycin,). Cells were cultured in a 37 °C, 5% CO_2_ environment for 10-14 days, after which mature MKs were enriched using size exclusion as described.

### Human megakaryocyte differentiation

CD34+ hematopoietic progenitor cells were isolated using the *EasySep™* Human CD34 Positive Selection Kit (STEMCELL Technologies, Cat. #100-1568), according to the manufacturer’s protocol. Cells were cultured in StemSpan medium (STEMCELL Technologies) supplemented with penicillin/streptomycin (100 U/mL, Sigma), L-glutamine (2 mM, Gibco),10 ng/mL TPO and 10 ng/mL interleukin-11 (Peprotech, UK) for 13 days.

### Naïve CD4□T cell isolation

Naïve CD4□T cells were isolated from mouse spleen single-cell suspensions using the EasySep™ Mouse Naïve CD4□T Cell Isolation Kit (STEMCELL Technologies, Cat. #19765), according to the manufacturer’s protocol.

### Murine ex vivo cultures

Bone marrow-derived megakaryocytes (MKs) were generated by culturing bone marrow progenitors in thrombopoietin (TPO)-containing medium for 10-14 days, as described above. Mature MKs were subsequently enriched by filter-based size exclusion. Prior to use in co-culture experiments, an aliquot of each MK preparation was analyzed by flow cytometry to assess purity and viability, and only preparations containing >90% viable MKs were used. Naïve CD4+ T cells were isolated and labelled with cell trace violet (ThermoFisher) according to manufacturer’s instructions. 4×10^4^ naïve CD4+ T cells were cultured with 1×10^4^ MKs and Ovalbumin peptides (OVA-323-339, 25 µg/mL, InvivoGen) for 4-5 days. Cells were cultured at 37 °C and 5% CO_2_ in U-bottom plates in a total volume of 200 μL of Dulbecco’s Modified Eagle’s Medium, GlutaMAX (DMEM GlutaMAX, Gibco) supplemented with 50 ng mL^-1^ TPO, 10% heat-inactivated Fetal Bovine Serum (FBS), sodium pyruvate, (1.5mM, Gibco), L-glutamine (2mM, Gibco), L-arginine (1x, Sigma), L-asparagine (1x, Sigma), penicillin/streptomycin (100 U/mL,Sigma), MEM non-essential amino acids (1x, ThermoFisher), and β-mercaptoethanol (57.2 uM, Sigma). As indicated, αMHC-II blocking antibody (10 µg/mL, M5/114.15.2, BioXCell) or a control IgG2b antibody (10 µg/mL, eB149/10H5, ThermoFisher) were added to cultures. At the completion of the co-culture period, cells were harvested and stained with both T cell and MK markers to simultaneously assess T cell responses and MK viability by flow cytometry.

### Ex Vivo iTreg cell induction

Sorted naïve CD4^+^CD62L^+^*Foxp3*^EGFP−^ T cells (4×10^4^) were stimulated with Dynabeads® Mouse T-Activator CD3/CD28 (11452D, Fisher Scientific) at a bead-to-cell ratio of 2:1 in the presence or absence of 1×10^4^ MKs. As indicated, anti-TGF-β (1,2,3) (10 μg/mL, MAB1835, R&D Systems) was added to the cultures.

### Intracellular Staining for Ex Vivo cultures

Following co-culture with megakaryocytes, T cells were stimulated with Cell Activation Cocktail (with Brefeldin A; BioLegend, Cat. No. 423303) for 4 hours to induce intracellular cytokines. Cells were subsequently stained for surface markers followed by fixation and permeabilization. Intracellular staining was performed for IFN-γ, IL-4, TNF-α, IL-17A, and expression of the lineage-associated transcription factors T-bet, GATA3, and RORγt.

### Cryosectioning and immunofluorescence staining

Tissues were fixed in 4% paraformaldehyde (PFA) in PBS overnight at 4 °C, then sequentially cryoprotected in 10%, 20%, and 30% sucrose solutions prepared in PBS (each for 24 hours at 4 °C). Dehydrated femurs were embedded in a water-soluble embedding medium and frozen at −20 °C. Using a tape transfer system (Kawamoto)(24), 10 µm cryosections were collected with a cryostat (Leica Biosystems). Sections were rehydrated in PBS for 20 min, blocked with 10% donkey serum and subsequently stained. Femoral cryosections were stained with CD105 (AF1320, R&D Systems), laminin (L9393, Sigma-Aldrich), LAP (AF-246-NA, R&D Systems), CD41 (133902, BioLegend) or CD42b (M-051-0, Emfret Analytics). Nuclei were stained using 4′,6-diamidino-2-phenylindole (DAPI). After washing in PBS with 0.1% Triton X-100, slides were mounted using Fluoroshield (F6182, Sigma-Aldrich) and imaged on a Zeiss LSM880 confocal microscope (20× objective). Whole femora were imaged on either a Lionheart automated microscope (BioTek) with a 4× objective, and MKs were quantified using Gen5 software. Mean fluorescent intensities (MFIs) were calculated and colocalization analysis was performed with Fiji Software (Version 2.14.0, NIH).

### Immunofluorescence staining of megakaryocytes and platelets

Primary bone marrow-derived or cultured MKs were isolated as described above and fixed in 4% PFA/PBS containing 0.1% Tween20 for 20 min at RT. Cells were spun down at 300 x g for 5 min and nonspecific antibody binding was blocked using 3% BSA/PBS. MKs were plated in 8-well chambers (155360, Nunc Lab-Tek), coated with an anti-mouse-CD31 antibody (102502, BioLegend) for 24 h and fixed with 4% paraformaldehyde. MKs and platelets were stained overnight using antibodies against LAP (AF246-NA, R&D Systems), a-tubulin (322588, ThermoFisher Scientific), proplatelet basic protein (PPBP), (PA5-47947, Invitrogen), CD42a (M-051-0, Emfret Analytics), Ovalbumin, Alexa Fluor™ 647 Conjugate (Thermo Fisher Scientific), nuclei were stained using DAPI. Cells were imaged at a Zeiss LSM880 confocal microscope (40x or 63x oil objective)

### Megakaryocyte ploidy by flow cytometry

Bone marrow single-cell suspensions were generated, and surface antigens’ staining was performed for 30 min on ice. Cells were then fixed, stained for SYTO™ 40 Blue (S1135, ThermoFisher Scientific) and analyzed according to the manufacturers protocol for True-Nuclear™ Transcription Factor Buffer Set (424401, Biolegend). MK DNA content was assessed by quantifying SYTO-40 distribution by flow cytometry.

### Live Cell Imaging of native megakaryocytes

Native MKs were isolated from the bone marrow of MHC II-eGFP mice as described. Cells were stained with SiR-DNA (1 µM, Cytoskeleton Inc.) for nuclear visualization and imaged live using a confocal fluorescence microscope (objective 63×, oil immersion). Brightfield and fluorescence channels were acquired simultaneously, and representative images were processed using ImageJ for background subtraction and contrast enhancement.

### Two-Photon intravital imaging of the calvaria

Intravital imaging of the bone marrow calvaria was performed as described (25). Briefly, a cranial window was generated by carefully thinning the parietal bone of anesthetized MHC II-EGFP reporter mice to allow optical access to the calvaria bone marrow. Mice were maintained under isoflurane anesthesia throughout the procedure, and body temperature was kept constant using a feedback-controlled heating pad. To visualize circulating platelets, mice received an intravenous injection of fluorescently conjugated anti-GPIbα antibody (0.1 µg/g body weight) immediately prior to imaging. Imaging was performed using a two-photon laser scanning microscope (excitation 920 nm, 20× water immersion objective). Image acquisition parameters (laser power, detector gain, and frame rate) were kept constant between experimental groups. For image analysis, 3D reconstructions were generated using Imaris (Bitplane), and colocalization between MHC II-EGFP+ cells and platelets was quantified using the Manders overlap coefficient on background-subtracted and thresholded images. Data were averaged from at least three independent fields per mouse, and four mice were analyzed.

### Megakaryocyte Sorting

Bone marrow was harvested from 10–12-week-old B6 mice and subjected to lineage depletion of lineage+ cells expressing CD3ε (clone 145-2C11), CD19 (clone 1D3), B220 (clone RA3-6B2), NK1.1 (clone PK136), CD11b (clone M1/70), CD11c (clone N418), Ly6G/C (clone RB6-8C5), and/or Ter119 (clone TER-119) using a biotin/streptavidin-based magnetic-activated cell sorting (MACS) approach on an autoMACS system (Miltenyi Biotec) prior to staining and flow cytometric sorting. Following lineage depletion, cells were stained for viability, DNA content, lineage markers, CD45, CD41, CD42d, and MHC II. Viable MKs were identified as (Live, Lineage^−^, CD45^+^, SYTO-40^+^, CD41^+^CD42d^+^) cells and were sorted into MHC II+ and MHC II− populations for RNA isolation using a FACSAria S6 (BD Biosciences).

### RNA Isolation

MKs (Live Lineage^−^ CD45^+^ SYTO-40^+^, CD41^+^CD42d^+^) were sorted from 10 to 12-week-old B6 mice and resuspended in Buffer RLT. For bulk RNA-seq, MKs were pooled from 4–5 mice to generate each library; four independent pooled libraries were analyzed per group. Individual mice contributed biological material to each pooled library but were not analyzed as independent RNA-seq replicates. RNA was extracted using RNeasy micro kit (QIAGEN), with 4 biological replicates per group.

### Bulk RNA Sequencing Analysis

Raw paired-end FASTQ files were processed using a standardized RNA-seq pipeline. Read quality was assessed with FastQC before and after trimming. Adapter removal and quality filtering were performed using fastp with default parameters. The GENCODE mouse reference (vM35, GRCm39) was downloaded and used to build a genome index with RSEM, enabling alignment through STAR. Gene-level quantification was performed with rsem-calculate-expression using the optio– --star, --paired-end, a– --strandedness reverse. Gene counts matrices were generated by combining the RSEM outputs across samples and filtering out genes with fewer than 10 total counts. Differential expression analysis between MHC-positive and MHC-negative groups was carried out using DESeq2 in R. Log2 fold changes were shrunk using ashr method, and genes with |log2 fold change| > 1 and adjusted p-value < 0.05 were considered differentially expressed. Functional enrichment analysis of up- and down-regulated genes was performed using clusterProfiler and msigdbr against the MSigDB C5 (Gene Ontology Biological Process) category.

### Integration of bulk RNA-seq with the Sun et al. single-cell MK atlas

An initial exploratory projection placed bulk samples on the reference UMAP by k-nearest-neighbour averaging of embedding coordinates. Because UMAP coordinates are non-metric and a bulk average does not correspond to a single embedded position, this exploratory view was superseded by the pseudobulk correlation and module-score analyses described above, which we report as the definitive integration. To relate the MHC II+ and MHC II-bulk transcriptomes to previously defined bone marrow megakaryocyte (MK) subsets, we integrated our data with the single-cell reference of Sun et al. (2). The reference was clustered at resolution 0.1, and the four clusters were annotated as platelet-producing, HSC-niche, inflammatory/immune, and actively cycling MKs using canonical subset markers (Pf4/Ppbp/Itga2b; Cxcl12/Mpl/Thbs1; H2-Aa/H2-Ab1/Cd74/Irf7; Mki67/Top2a/Birc5, respectively). The assignment was confirmed by marker expression. For each subset we generated pseudobulk profiles by summing raw UMI counts across the constituent cells. To estimate within-subset variability, cells in each subset were randomly partitioned into three pseudo-replicates. A unified gene panel (56 genes) was assembled from two sources: subset-defining signature genes derived directly from the reference by differential expression (FindAllMarkers, Wilcoxon, adjusted P < 0.05; top genes per subset by log2 fold change), and a curated MHC class II and co-stimulatory module (H2-Ab1, H2-Aa, H2-Eb1, H2-DMa, H2-DMb1, Cd74, Ciita, Cd80, Cd86, Cd40, B2m, Tap1/2, Psmb8/9). Pseudobulk and bulk counts were normalized to log2 counts-per-million (edgeR), and the panel was visualized as row z-scored heatmaps for the reference subsets and for the bulk MHC II+/- samples in parallel.

To map each bulk sample to the reference, Spearman correlations were computed between every bulk replicate and each subset pseudobulk across the 2,000 shared highly variable genes, and each replicate was assigned to its maximally correlated subset. The specificity of this mapping was validated by passing the subset pseudo-replicates back through the identical correlation pipeline, which recovered the correct subset of origin in all 12 of 12 subset pseudo-replicates. Finally, MHC II+ and MHC II-bulk signatures (DESeq2 DEGs, absolute log2FC > 1, adjusted P < 0.05; 518 and 235 genes, respectively) were scored across all reference cells with AddModuleScore and compared across subsets. Analyses used R v4.6.0 with Seurat v5.5.1, SeuratObject v5.4.0, DESeq2 v1.52.0, edgeR v4.10.1, limma v3.68.4, pheatmap v1.0.13, and ggplot2 v4.0.3.

### Transcription Factor Activity Inference

Transcription factor (TF) activity was inferred using the VIPER algorithm implemented via the decoupleR package with regulons obtained from DoRothEA (mus musculus version). Regulons with confidence levels A–C were included. Variance-stabilized expression values served as input, and activity scores were calculated for each TF in each sample. Mean activity per TF per group was calculated and differential activity (Δ Activity) was defined as the difference between MHC-positive and MHC-negative mean scores. TFs were ranked by the absolute magnitude of Δ Activity, and the top TFs (n = 35) were selected for visualization.

### Statistics

Statistical analyses are reported as mean ± SD or ± SEM. A Shapiro-Wilk test was used to determine normal versus abnormal distributions, and all continuous variables were tested for mean differences. Depending on the spread of variable, both nonparametric Mann-Whitney *U* test, ANOVA, Kruskal-Wallis, and Wilcoxon tests and parametric 2-tailed Student’s *t* test and 1- or 2-way ANOVA were used. For ANOVA, Tukey’s post hoc test was used to compare individual groups. Data were analyzed and visualized using R (v.4.5.1).

## Supplementary Materials

Supplementary Figures S1 to S7

Supplementary Table 1-2

Supplementary Movies 1, 2, and 3

## Acknowledgments

We would like to acknowledge Dr. Robert Welner, Dr. Robert Signer, and Dr. Jarrod Dudakov for their helpful and insightful conversations.

## Funding

National Institutes of Health National Institute of Diabetes and Digestive and Kidney Diseases R01DK139341 (KRM)

National Institutes of Health National Institute of Diabetes and Digestive and Kidney Diseases R21DK142111 (KRM)

National Institutes of Health National Heart, Lung, and Blood Institute R01HL151494 (KRM)

National Institutes of Health National Heart, Lung, and Blood Institute K99HL175037 (MNB)

National Institutes of Health National Heart, Lung, and Blood Institute R35HL161175 (JEI)

American Society of Hematology Restart Award (MNB)

American Society of Hematology Scholar Award (VC)

American Society of Hematology Treating Fairly Award (KRM)

American Heart Association 24IPA1274573 (KRM)

American Heart Association 23POST1011433 (MNB)

American Heart Association 25DIVSUP1476419 (VC)

VGS is an investigator of the Howard Hughes Medical Institute.

## Author contributions

Conceptualization: VC, KRM

Experimental Design: VC, KRM

Experimentation: VC, KGW, EC, ICB, DHL, DG, CW, JM, MC

Data Analysis: VC, MF, JM, MAB

Statistics: VC

Funding acquisition: VC, UvA, VGS, JEI, KRM

Supervision: JEI, KRM

Writing – original draft: VC, KRM

Writing – editing: VC, KRM, MAB, EC, MNB, PH

## Data and materials availability

All data, code, and materials used in the analysis will be available upon reasonable request made to the corresponding author.

