## Supplementary Figures for "MHC II-expressing bone marrow megakaryocytes are noncanonical antigen presenting cells and activate CD4^+^ T cells ex vivo"

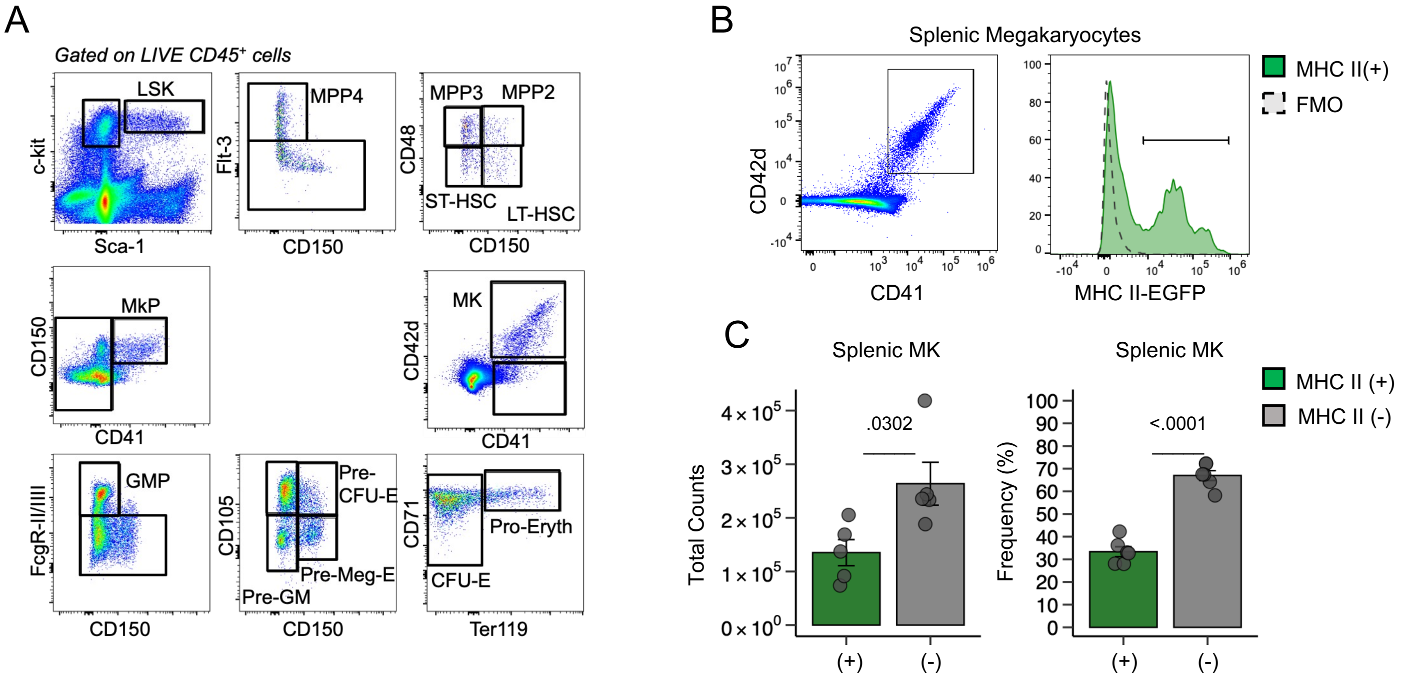


**SUPPLEMENTARY FIGURE 1**

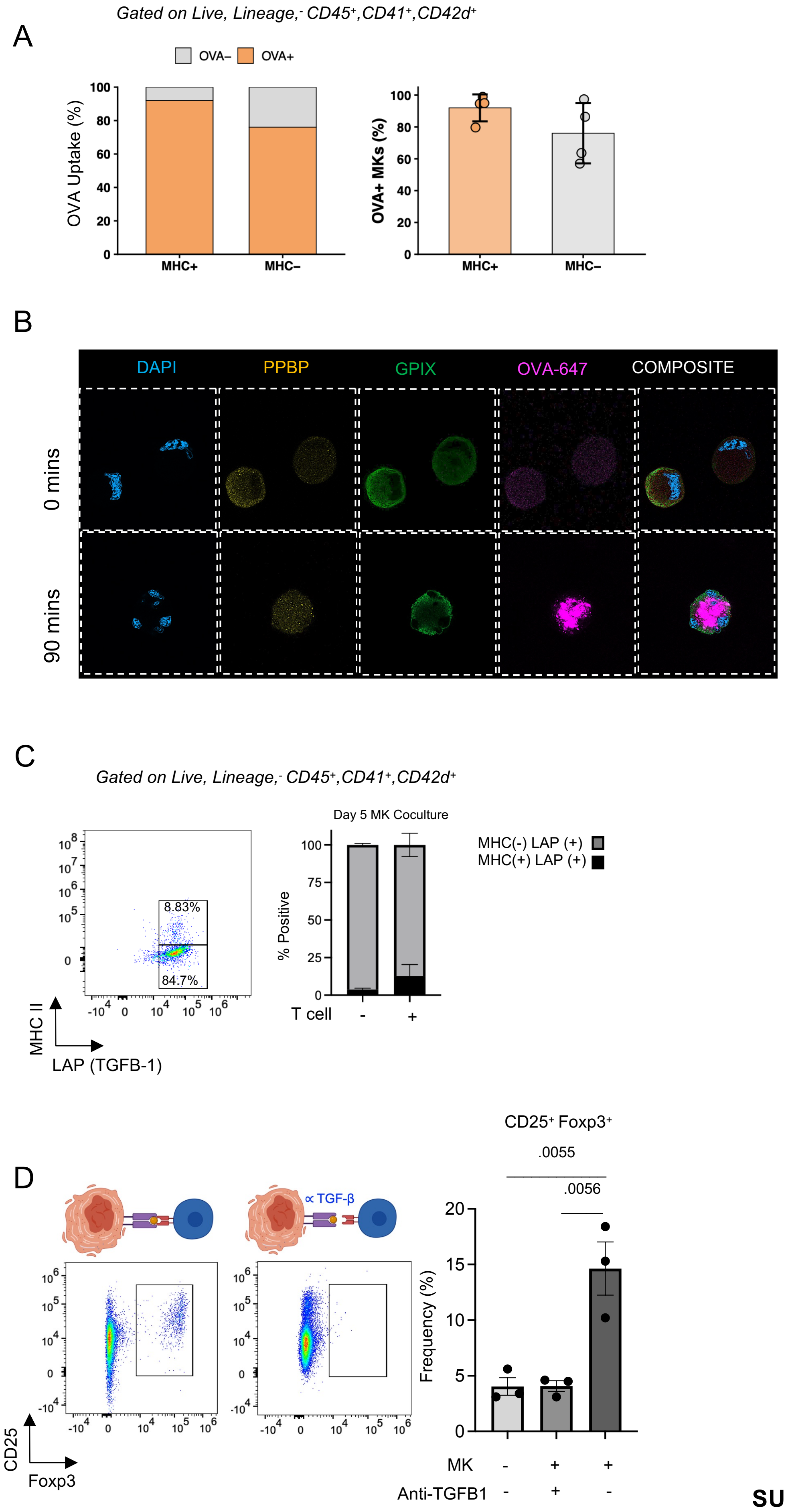


**SUPPLEMENTARY FIGURE 2**

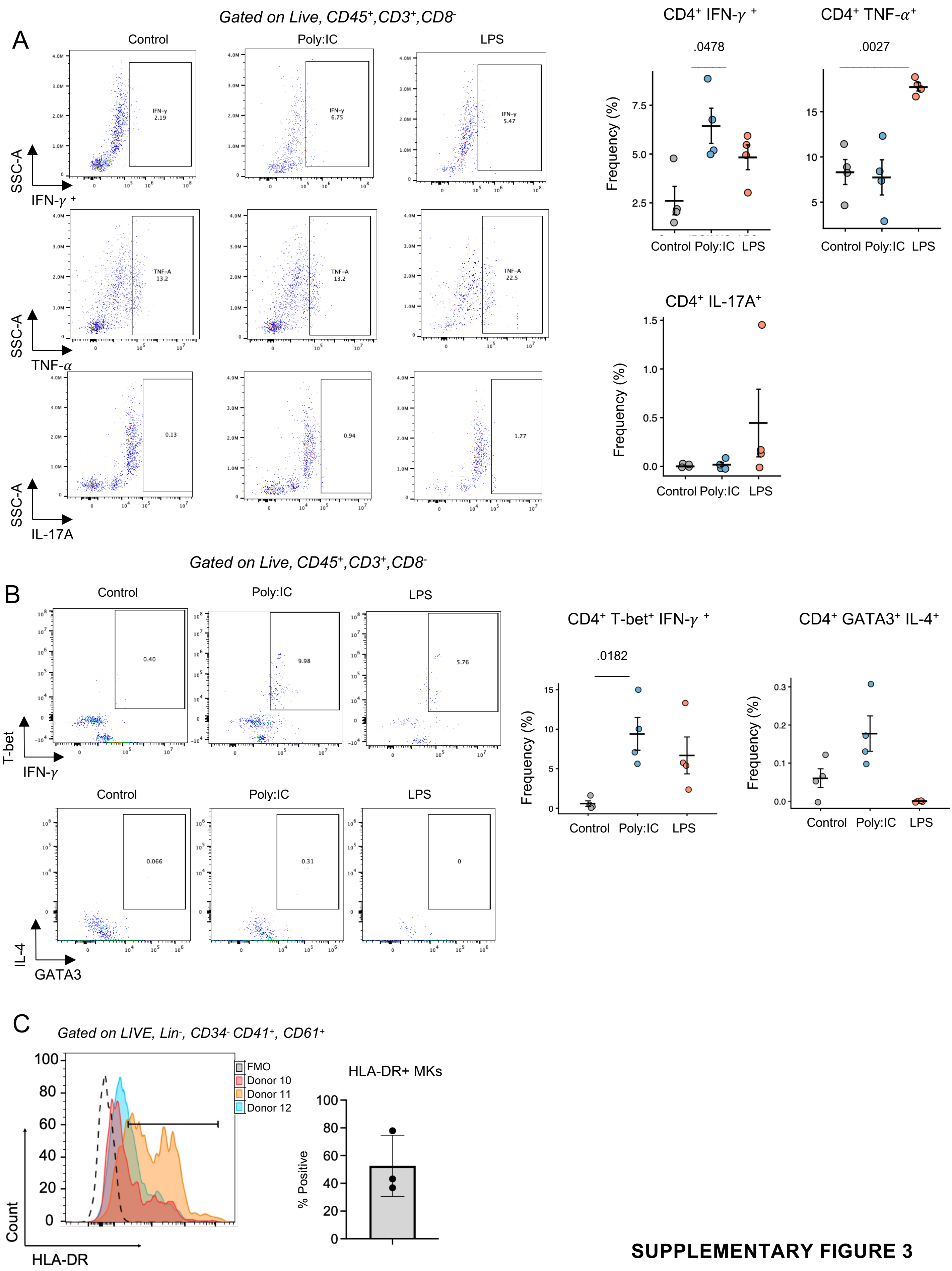


**SUPPLEMENTARY FIGURE 3**

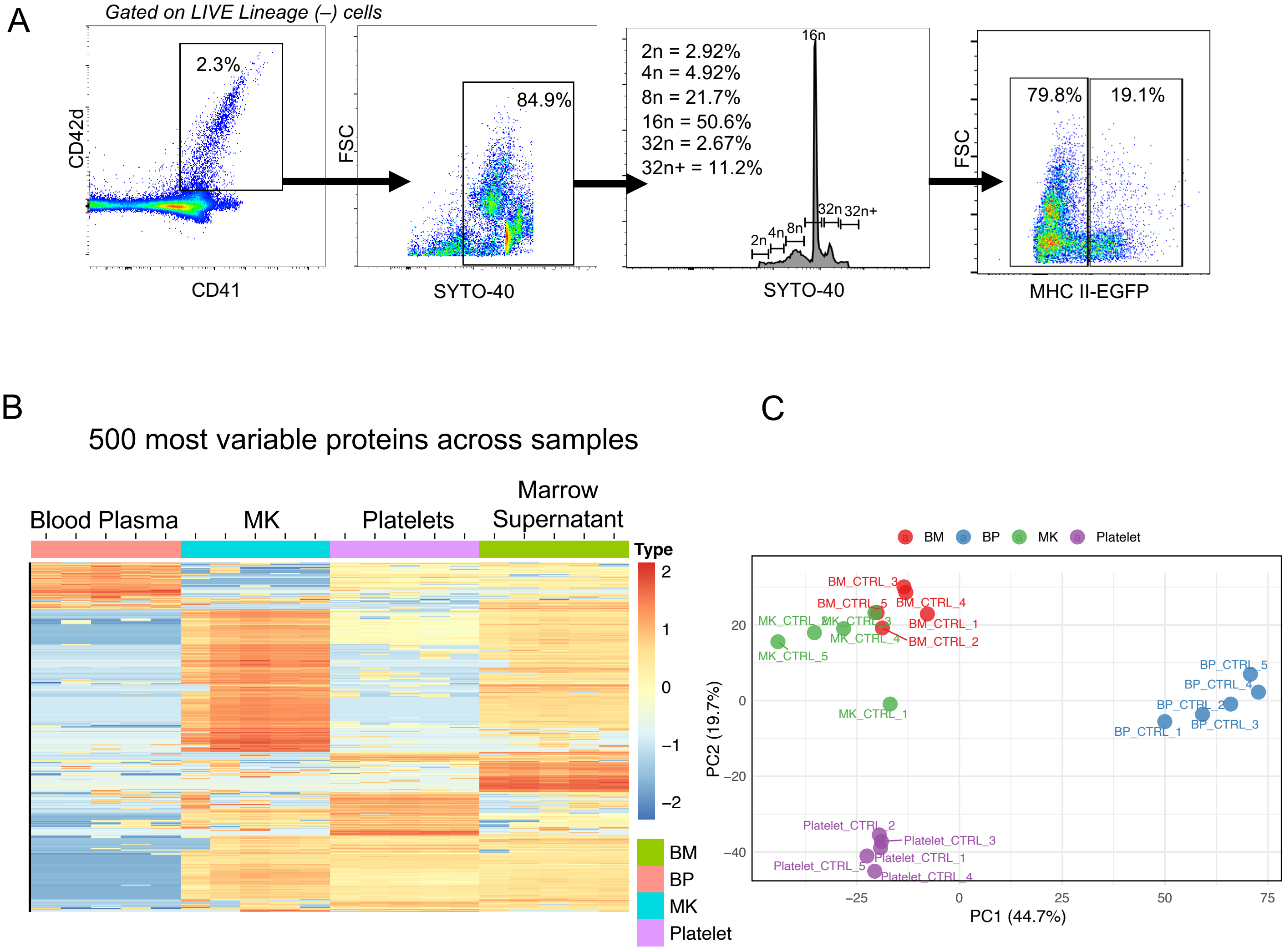


**SUPPLEMENTARY FIGURE 4**

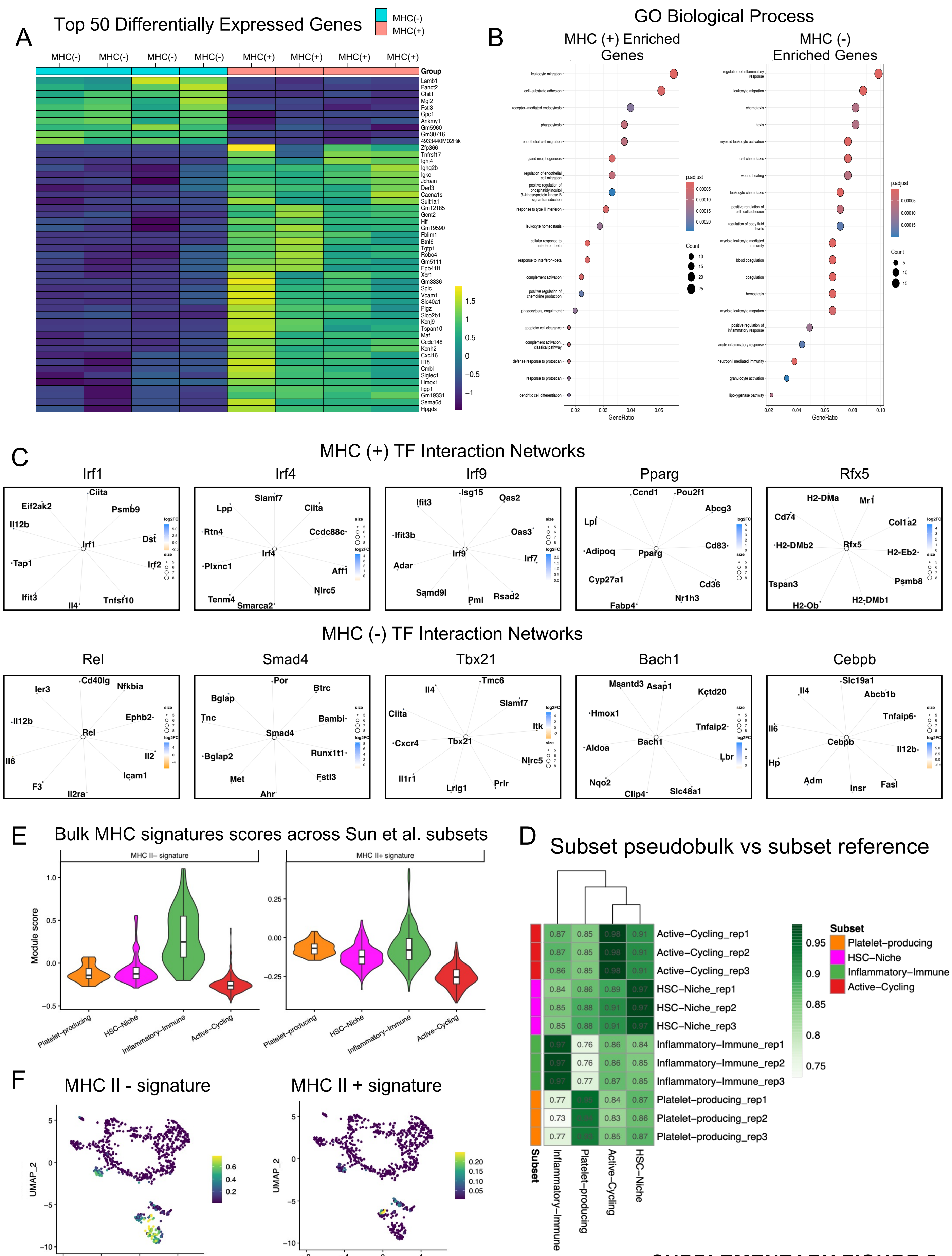


SUPPLEMENTARY FIGURE 5

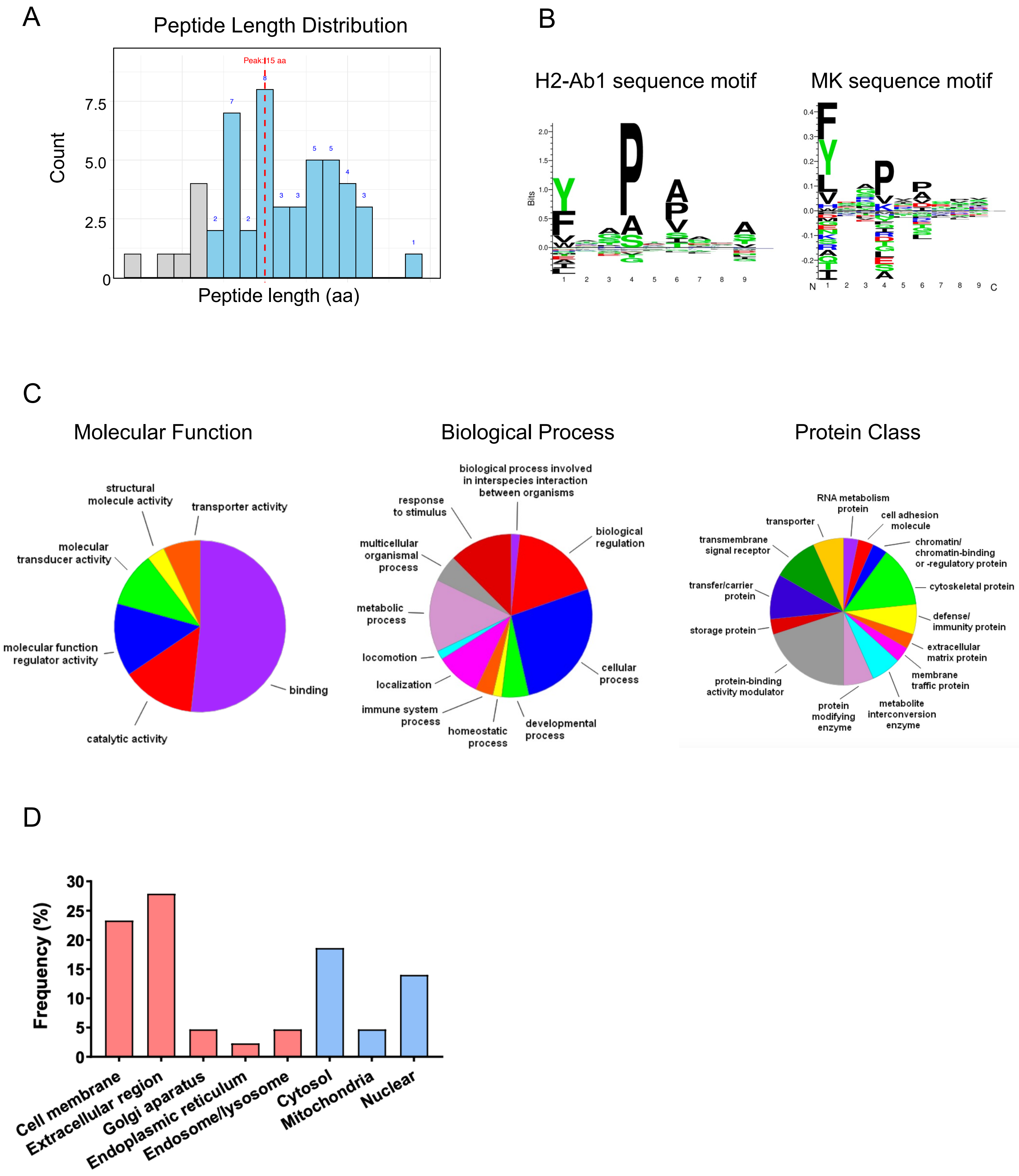


SUPPLEMENTARY FIGURE 6

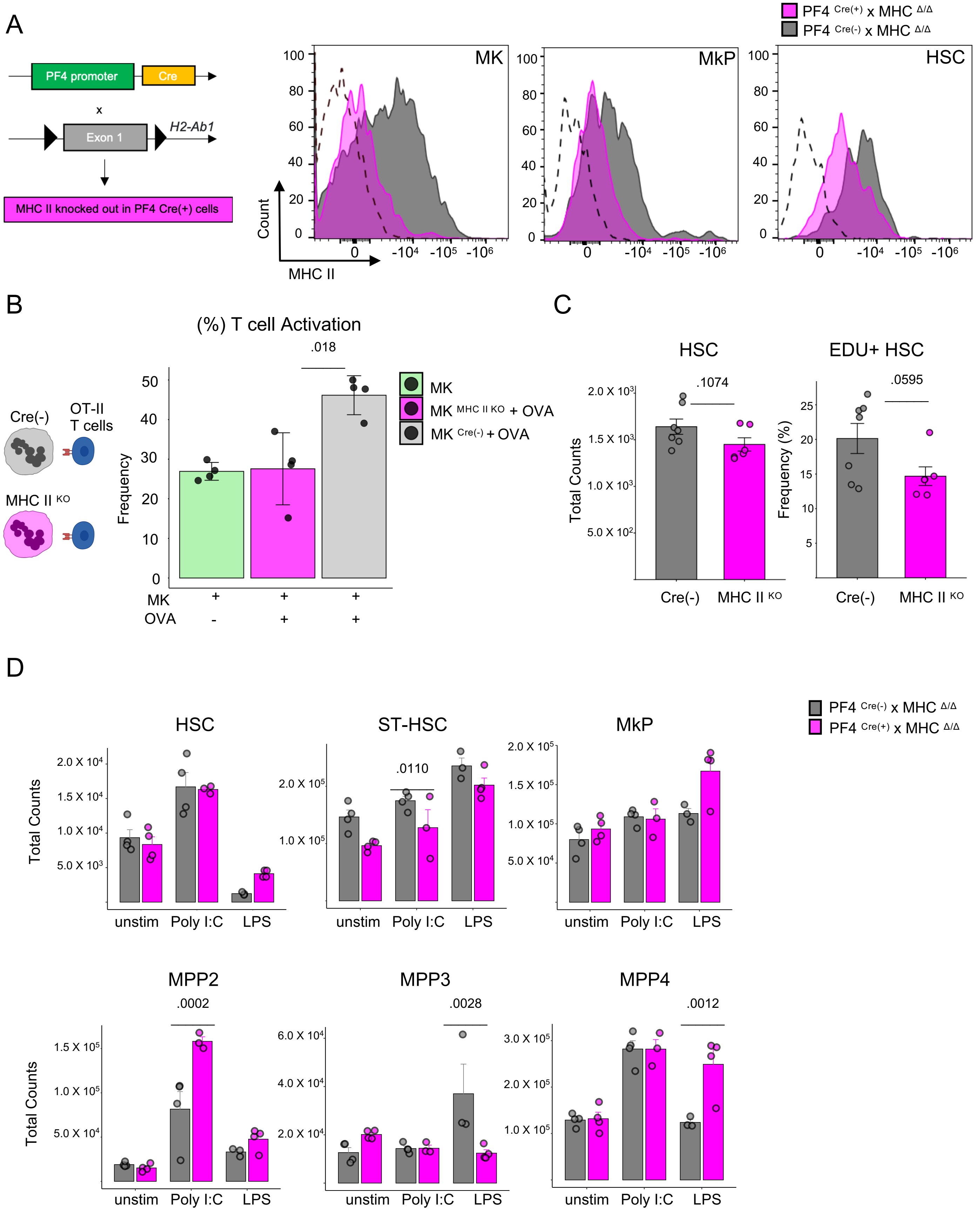


**SUPPLEMENTARY FIGURE 7**
