## Supplementary Figure Legends for "MHC II-expressing bone marrow megakaryocytes are noncanonical antigen presenting cells and activate CD4^+^ T cells ex vivo"

**Fig. S1. MK MHC II is differentially expressed in bone marrow and spleen.**

(**A**) Gating strategy of bone marrow progenitor populations.

(**B**) Representative plots and histogram of MHC II expression in splenic MKs.

(**C**) Absolute counts and frequencies of MHC II+ and MHC II- splenic MKs; *n* = 5 animals. Data are shown as mean ± SD; graphs represent data from at least 3 independent experiments. Statistics performed with unpaired 2-tailed Student *t* test.

**Fig. S2. MHC II+ MKs interact with T cells.**

(**A**) Flow cytometry quantification of OVA uptake in bone marrow MKs; OVA+ (orange), OVA-(grey); right, and frequency of OVA+ cells in either MHC II+ or MHC II- MKs; n=4 animals.

(**B**) Representative images of OVA uptake in bone marrow MKs; DAPI (blue), PPBP (yellow), OVA-647 (magenta), GPIX (green).

(**C**) Representative plots and quantification of intracellular LAP in MKs in the presence or absence of T cells n=4 animals/ condition.

(**D**) Representative plots and quantification (Frequencies) of CD25+ Foxp3+ cells in MK and T cell co-cultures of T cells n=3 animals/ condition. Data are shown as mean ± SEM; graphs represent data from at least 3 independent experiments. Statistics performed with unpaired 2-tailed Student *t* test (B) or and 1-way ANOVA with Tukey’s multiple comparisons test (A, C).

**Fig. S3. MHC II+ MKs activate T cells ex vivo.**

(**A**) Representative plots and quantification (Frequencies) of IFN-y^+^, TNF-a^+^ and IL-17^+^ CD4 T cells in MK and T cell co-cultures (+/- MK stimulation); n=4 animals/ condition.

(**B**) Representative plots and quantification (Frequencies) of (Th1: T-bet^+^, IFN-y^+^), and (Th2: GATA3^+^ IL-4^+^) CD4 T cells in MK and T cell co-cultures (+/- MK stimulation); n=4 animals/ condition.

(**C**) Representative histogram of HLA-DR expression in MKs isolated from human PBMCs.

Data are shown as mean ± SEM; graphs represent data from at least 2 independent experiments. Statistics performed with 1-way ANOVA with Tukey’s multiple comparisons test (A-B). Th2 frequencies were analyzed using a Kruskal–Wallis test followed by Dunn's multiple-comparisons test with Holm correction.

**Fig. S4. MHC II is not present on platelets.**

(**A**) Gating strategy of bone marrow MK DNA content.

(**B**) Heatmap of 500 differentially expressed proteins across MKs, blood plasma, bone marrow fluid, and platelets; *n* = 4 animals.

(**C**) PCA plot of sample distribution; *n* = 4 animals.

**Fig. S5. MHC II- and MHC**+ **MKs have distinct transcriptional programs.**

(**A**) Heatmap of top 50 differentially expressed genes between MHC II- and MHC II+ MKs.

(**B**) Dot plot of gene set enrichment analysis of the GO biological pathways associated with MHC II- and MHC II+ MKs.

(**C**) Inferred transcription factor (TF) regulatory networks highlight distinct transcriptional programs in MHC II+ MKs (Irf4, Rfx5, Irf9, Pparg, Irf1) (top) and MHC II- MKs (Smad4, Tbx21, Bach1, Rel, Cebpb) (bottom), with node color indicating differential gene expression (log₂FC) and edges representing predicted TF-target interactions.

**(D)** MHC II+ and MHC II- signature scoring across single cells and the pseudobulk validation control (12 of 12 correct).

**(E)** Signature Feature plot of MHC II+ and MHC II- MKs

**(F)** Violin Plots of Bulk MHC II+ and MHC II- signature scores across the Sun. et al. subsets.

**Fig. S6. Immunopeptidomics reveals exogenous peptides on MK MHC II.**

(**A**) Length distribution of peptides isolated from MK MHC II molecules.

(**B**) Clustering results for the peptide dataset; the corresponding sequence motif is shown as sequence logos (left). H2-Ab1motif available in NetMHCIIpan v4.1 database obtained from integration and motif deconvolution of mass spectrometry MHC eluted ligand data.

(**C**) Gene ontology analysis: Frequency of proteins associated to different molecular functions, biological processes, and protein class, based on the Gene Ontology database annotations for parental proteins of MHC II-isolated peptides

(**D**) Degradative pathway of the MHC II-associated peptide parental proteins. Endo-lysosomal degradation pathway (red) includes proteins from cell membrane, extracellular region, Golgi apparatus, endoplasmic reticulum, and endosome/lysosome. Cytosolic, mitochondrial, and nuclear proteins were included as the cytosolic pathway (blue) of degradation

**Fig. S7. MK-specific MHC II deletion alters HSPC response to LPS and Poly (I:C) administratio**n

(**A**) Schematic of PF4-targeted MHC II deletion and representative histograms of MHC II expression in MKs, MkPs, and HSCs.

(**B**) Quantification of T cell activation in the presence and absence of OVA in MHC^KO^ and Cre ^(-)^ mice; *n* =4 mice/ condition.

(**C**) Total counts of HSCs (right) and frequency of EdU+ HSCs in MHC^KO^ and Cre ^(-)^ mice (left); *n* =5 mice/ genotype.

(**D**) Total counts of HSCs, ST-HSCs, MkPs, MPP2s, MMP3s, and MPP4s 24 hours after LPS or Poly (I:C) administration; *n* = 4 mice/ genotype. Data are shown as mean ± SD; graphs represent data from at least 3 independent experiments. Statistics performed with unpaired 2-tailed Student t test (C-D) and 1-way ANOVA (B) and 2-way ANOVA (D) with Tukey’ multiple comparisons test at 95.00% CI.
